# Impact of exogenous aminoacyl-tRNA synthetase and tRNA on temperature sensitivity in *Escherichia coli*

**DOI:** 10.1101/2024.05.02.592135

**Authors:** Jongdoo Choi, Jiyeun Ahn, Jieun Bae, Moonsang Yoon, Hwayoung Yun, Minseob Koh

## Abstract

Genetic code expansion (GCE) is a powerful strategy that expands the genetic code of an organism for incorporating non-canonical amino acids (ncAAs) into proteins using engineered tRNAs and aminoacyl-tRNA synthetases (aaRSs). While GCE has opened up new possibilities for synthetic biology, little is known about the potential side effects of exogenous aaRS/tRNA pairs. In this study, we investigated the impact of exogenous aaRS and amber suppressor tRNA on gene expression in *Escherichia coli*. We discovered that in DH10β Δ*cyaA*, transformed with the F1RP/F2P two-hybrid system, high consumption rate of cellular ATP by exogenous aaRS/tRNA at elevated temperatures induces temperature sensitivity in the expression of genes regulated by the catabolite activator protein. We harnessed this temperature sensitivity to create a novel biological AND gate in *E. coli*, responsive to both *p*-benzoylphenylalanine (BzF) and low temperature, using a BzF-dependent variant of *E. coli* chorismate mutase and split subunits of *Bordetella pertussis* adenylate cyclase. Our study provides new insights into the unexpected effects of exogenous aaRS/tRNA pairs and offers a new approach for constructing a biological logic gate.

## Introduction

The genetic code expansion (GCE) strategy has emerged as a powerful tool for site-specifically incorporating non-canonical amino acids (ncAAs) into proteins, unlocking new avenues in the field of synthetic biology and protein engineering.^1–3^ By expanding the genetic code beyond the conventional 20 amino acids, GCE empowers researchers to embed unique functionalities into proteins, paving the way for myriad applications. For example, through GCE, enzymes can be fine-tuned for superior catalytic efficiency^4^ or heightened stability^5, 6^ by incorporating ncAAs. Moreover, the strategic use of ncAAs with bioorthogonal chemistry functionalities has facilitated the direct visualization and labeling of biomolecules in live cells,^7^ while also optimizing the synthesis of antibody–drug conjugates to produce homogeneous batches.^1^ Beyond these applications, GCE offers the potential to engineer synthetic organisms endowed with ncAA-driven novel functionalities or characteristics.^8–13^

The GCE strategy requires expressing exogenous aminoacyl-tRNA synthetase (aaRS) and transfer RNA (tRNA) pair to suppress a blank codon.^3, 14^ While GCE presents immense potential, the overexpression of exogenous aaRS and tRNA has raised concerns about potential side effects on cellular processes.^15^ The possible effects include disruption of ribosomal activity by the engineered tRNA,^16^ destruction of innate protein activity through C-terminal extension,^17^ potential cellular burden due to the production of truncated proteins,^18^ and dysregulation in stress response pathways.^19^ Although these effects appear to be marginal, considering the numerous successful applications of GCE across various organisms including bacteria, mammalian cells, and even animals,^3^ the impact can become substantial under extreme conditions. This is especially true in knockout host organisms where metabolic systems might be significantly altered or compromised.

The *cyaA* gene encodes adenylate cyclase, responsible for converting adenosine triphosphate (ATP) into adenosine 3’,5’-monophosphate (cyclic AMP, cAMP), and is associated with carbon metabolism in *E. coli*.^20^ The binding of cAMP to its receptor protein (cAMP Receptor Protein, CRP) triggers a conformational change in the CRP, allowing it to bind to its cognate promoters and drive downstream gene expression.^21^ Given that approximately 380 promoters in *E. coli* are known to be dependent on the cAMP–CRP,^22^ a knockout of *cyaA* could be detrimental to normal growth. ATP biosynthesis is also largely dependent on the cAMP– CRP system.^22^ Considering that the ligation of tRNA to its paired amino acid is catalyzed by its cognate aaRS and is absolutely dependent on ATP,^23^ the function of the overexpressed aaRS/tRNA may affect the cAMP-dependent gene expression in the *cyaA* knockout strain, or vice versa.

During our efforts to develop a novel protein–protein interaction (PPI) module that incorporates an ncAA at the PPI interface, we serendipitously discovered the GCE-caused temperature sensitivity in *cyaA* knockout *E. coli*. We used a bacterial adenylate cyclase two-hybrid (BACTH) system^24^ by constructing fusions of a *p*-benzoylphenylalanine (BzF) dependent variant of *E. coli* chorismate mutase (ECB)^25^ with split subunits (T18 and T25) of *Bordetella pertussis* adenylate cyclase. The requirement of the BzFRS/tRNA_CUA_ pair^26^ for incorporating BzF to the ECB interface led us to discover ATP consumption by these components significantly affects the temperature sensitivity of cAMP-dependent gene expression in *cyaA* knockout *E. coli*. In our experiments using DH10β Δ*cyaA* cells harboring pUltra-BzF plasmid,^25^ genes under the control of the *lac* promoter exhibited expression levels inversely proportional to the temperature increase (Fig. 1).

**Fig. 1.**
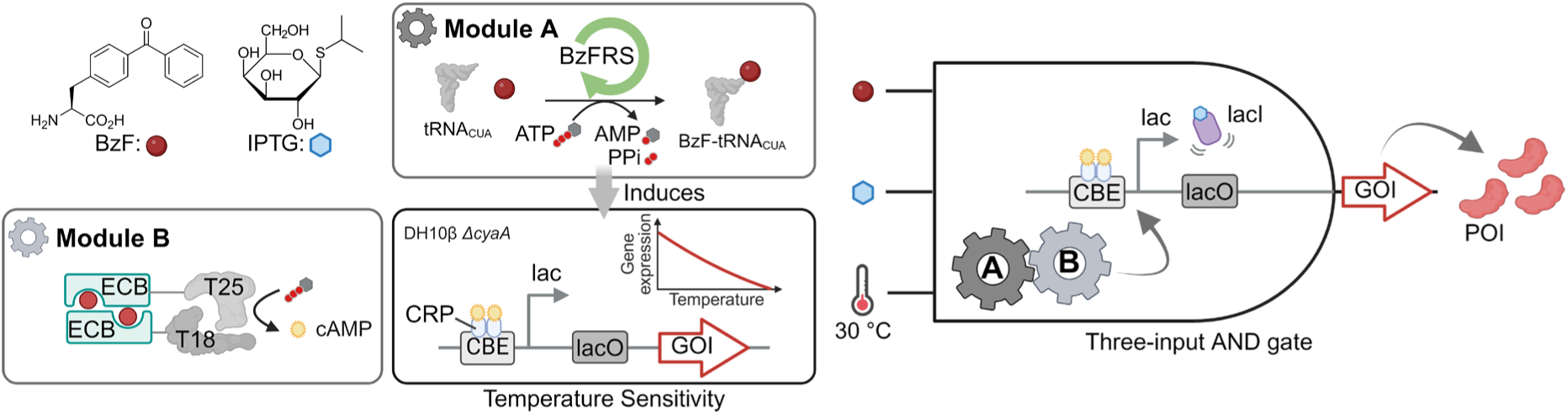
Schematic illustration depicting the temperature sensitivity induced by exogenous aminoacyl-tRNA synthetase and tRNA in *cyaA* knockout *E. coli*, along with a three-input biological AND gate utilizing BzF, IPTG, and low temperature (30 °C) as inputs. BzF, *p*-benzoylphenylalanine; BzFRS, *p*-benzoylphenylalanyl-tRNA ligase; ECB, BzF dependent variant of *E. coli* chorismate mutase; T25 and T18, split subunits of *Bordetella pertussis* adenylate cyclase; lacI, *lac* repressor; CRP, cAMP receptor protein; CBE, CRP binding element; lacO, *lac* operator; GOI, gene of interest; and POI, protein of interest.

Taking advantage of the unexpected temperature sensitivity caused by the exogenous aaRS/tRNA, we developed a novel biological AND gate. Because cAMP production is crucial for the expression of gene of interest (GOI), any inputs stimulating cAMP production can be utilized for GOI expression. The ncAA BzF can be one such input, given its essential role in the PPI module (ECB) for cAMP production, along with the requirement of a low-temperature condition. The system displayed a significant dynamic range in the regulation of target gene expression. Specifically, under permissive conditions with both inputs (0.5 mM BzF and 25 °C) present, we observed a maximum 86-fold amplification compared to non-permissive conditions lacking one or both inputs (Supplementary Fig. 1). We also have demonstrated a three-input AND gate system utilizing BzF, low temperature (30 °C), and Isopropyl β-D-1-thiogalactopyranoside (IPTG) as three distinct inputs (Fig. 1). This system can be used to design novel biologic gates that are distinct from conventional versions.^27, 28^

## Results

### Design of a bacterial two-hybrid system

In our study, we engineered a BACTH system^24^ that is built around ECM, a wild type *E. coli* chorismate mutase. Our design strategy was based on the homodimeric nature of ECM.^29^ The ECM-BACTH system includes two key plasmids: F1RP (Fusion 1 Reporter Plasmid), encoding a T25-ECM fusion protein and a reporter gene, and F2P (Fusion 2 Plasmid), encoding an ECM-T18 fusion protein (Supplementary Fig. 2A). The functionality of our system relies on ECM dimerization, which triggers the dimerization of T18-T25 and subsequent cAMP generation. Reporter genes in this system are controlled by the *lac* promoter, modulating their expression through the cAMP–CRP.^22^

The ability of our system to modulate gene expression in response to ECM dimerization was tested in a growth experiment. In this experiment, we introduced the gene encoding chloramphenicol acetyl transferase (CAT) into the F1RP plasmid, creating F1RP-CAT (see Supplementary Table 1 for detailed plasmid information). This was then transformed into DH10β Δ*cyaA* cells along with F2P. The survival of the resultant DH10β Δ*cyaA*/F1RP-CAT/F2P cells in chloramphenicol (Cm) media was confirmed, demonstrating the ability of the ECM dimer to function as a PPI module for gene regulation (Supplementary Fig. 2B). Building upon the ncAA-dependent ECM variant (ECB) presented in previous research,^25^ and informed by our subsequent experiments, we gained our confidence in the ECM-BACTH system as a robust platform for ncAA-dependent gene regulation.

### Exogenous aaRS/tRNA confers temperature sensitivity in cAMP dependent gene expression in ECM-BACTH

Given the inherent cAMP biosynthesis deficiency in the DH10β Δ*cyaA* strain, which may lead to functional complications, we postulated that this strain would exhibit a heightened sensitivity to external stimuli. Therefore, prior to developing the ncAA-dependent version of ECM-BACTH system, it was essential to assess any unintended effects stemming from the expression of BzFRS and tRNA_CUA_, which are key components in the GCE strategy. To evaluate the impact of these exogenous factors, we incorporated the SP plasmid (pUltra-BzF),^25^ which expresses BzFRS/tRNA_CUA_, into our ECM-BACTH system and monitored the expression of the *lac* promoter-controlled GOI.

An unexpected observation that emerged during our initial experiments was the pronounced sensitivity of gene expression to temperature changes. This was most notable during a growth experiment with DH10β Δ*cyaA*/F1RP-CAT/F2P/SP cells incubated on LB-agar at either 30 °C or 37 °C. We found that the presence of BzFRS/tRNA_CUA_ diminished cell survival in Cm media, an effect that was significantly amplified with the addition of BzF (0.5 mM) at the higher temperature (37 °C), leading to total cell death in high-Cm conditions (150 μg/mL) (Fig. 2B).

**Fig. 2.**
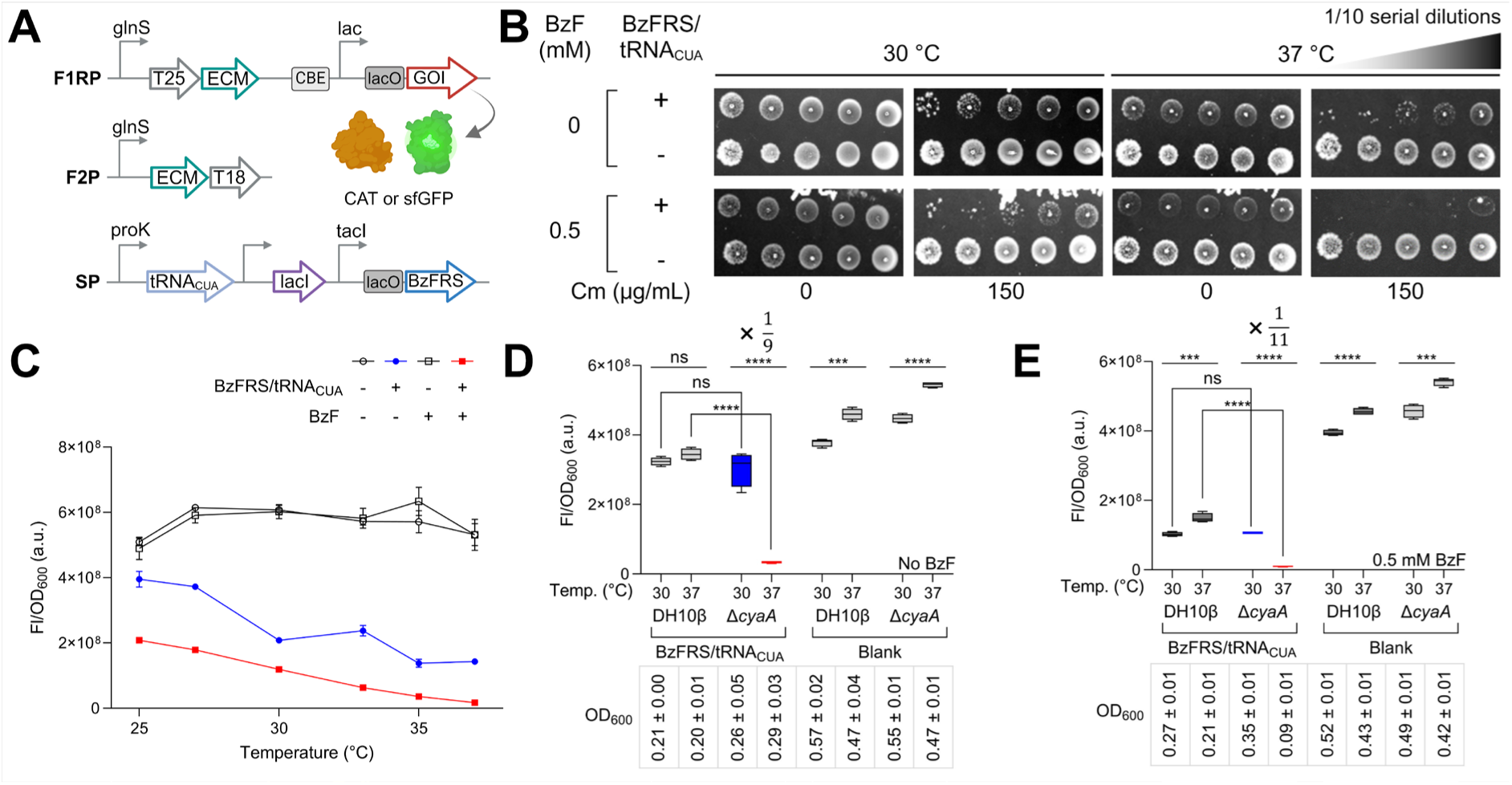
Impact of exogenous BzFRS/tRNA_CUA_ on temperature sensitivity in gene expression in DH10β Δ*cyaA*. An *E. coli* chorismate mutase (ECM) mediated bacterial adenylate cyclase two-hybrid (ECM-BACTH) system complements the cyclic AMP (cAMP) deficiency, and all the tested genes are under the control of the cAMP-dependent *lac* promoter. **A** The scheme of the plasmids. ‘F1RP’ represents the T25-ECM fusion plasmid containing the reporter gene, ‘F2P’ refers to the ECM-T18 fusion plasmid, and ‘SP’ means pUltra-BzF. CAT represents chloramphenicol acetyltransferase and sfGFP refers super-folder green fluorescent protein. **B** Dilution spot images of the DH10β Δ*cyaA*/F1RP-CAT/F2P harboring SP or SP^null^ on LB-agar, both in the absence and presence of chloramphenicol (Cm, 150 μg/mL), incubated under specified temperatures for 72 h. **C** The normalized sfGFP fluorescence signals were plotted over temperatures (6 points; 25 °C, 27 °C, 30 °C, 33 °C, 35 °C, and 37 °C) after incubating DH10β Δ*cyaA*/F1RP-GFP/F2P cells containing SP or SP^null^ in the specified conditions for 24 h. **D**, **E** DH10β and DH10β Δ*cyaA*/F1RP-GFP/F2P cells harboring SP or SP^null^ were incubated in LB broth in the absence (**D**) and presence (**E**) of BzF (0.5 mM) under specified temperatures for 24 h and the expression levels of the sfGFP were quantified by fluorescence reading (Ex/Em: 485 nm/525 nm). Blank plasmids were constructed by deleting BzFRS/tRNA_CUA_ (SP^null^); n = 4 (**C**, **D**, and **E**). Error bars stand for standard deviation (**C**); box limits indicate the interquartile range; whiskers represent the range from minimum to maximum; and the center line denotes the median (**D** and **E**). Fl, fluorescence; OD_600_, optical density at 600 nm; ns (not significant), *** P < 0.001, **** P < 0.0001 by Student’s t-test; and variability in OD_600_ values is represented by the standard deviation.

Motivated by the intriguing temperature sensitivity, we conducted sfGFP expression measurements across a range of temperatures, both with and without BzF (Fig. 2C and Supplementary Fig. 1A). Remarkably, in the presence of BzFRS and tRNA_CUA_, we observed a consistent suppression of sfGFP expression across the entire temperature range, which intensified at higher temperatures, suggesting that BzFRS and tRNA_CUA_ negatively impact GOI expression in the ECM-BACTH, especially at higher temperatures. Notably, the enhanced expression at the lower temperature (30 °C) was specific to *cyaA* knockout DH10β expressing BzFRS/tRNA_CUA_. Furthermore, the difference in expression levels between the two temperatures (30 °C and 37 °C) became even more pronounced in the presence of BzF (0.5 mM), exhibiting a decrease by a factor of 1/11 with BzF and 1/9 without BzF (Fig. 2D and 2E).

To investigate whether BzFRS and tRNA_CUA_ contribute individually or collectively to this temperature sensitivity, we compared the effects of various control plasmids (SP-aaRS^null^, SP-tRNA^null^, and SP^null^ plasmid) on sfGFP expression. These tests were performed on DH10β Δ*cyaA*/F1RP-GFP/F2P cells at 30 °C and 37 °C in the presence and absence of BzF. Intriguingly, we found that not only the concurrent expression of BzFRS and tRNA_CUA_, but also their individual expression, led to a higher sfGFP expression at 30 °C compared to 37 °C. The SP, SP-aaRS^null^, and SP-tRNA^null^ led to 7.4-fold, 1.4-fold, and 2.5-fold reductions in sfGFP fluorescence, respectively, in the absence of BzF at the higher temperature. Notably, the SP-aaRS^null^ condition was unaffected by BzF. In contrast, BzF presence consistently reduced sfGFP fluorescence for SP and SP-tRNA^null^, with increased deviation to 7.6-fold and 9.5-fold for SP and SP-tRNA^null^, respectively (Fig. 3). To address this, we proposed that the observed temperature sensitivity may be attributed to ATP depletion occurring during catalysis involving the exogenous aaRS and tRNA. We also consider unintended cross-reactions between the evolved archaeal aaRS/tRNA and the native *E. coli* aaRS/tRNAs, as well as with natural amino acids, to be one mechanism underlying the depletion of ATP. In this context, we confirmed tyrosine incorporation at the amber codon site of sfGFP in the absence of BzF (Supplementary Fig. 3 and Supplementary Table 2). Notably, under the conditions of exogenous tRNA_CUA_ alone, we observed the incorporation of leucine, glutamate, proline, and tyrosine at the amber codon site, as assessed by mass spectrometry. This occurred regardless of the presence of BzF (Supplementary Fig. 4 and Supplementary Table 2). This suggests that the evolved tRNA_CUA_ may exhibit some inherent promiscuity, allowing it to accept certain natural amino acids in the absence of its cognate aaRS (BzFRS). This aligns with previous findings on the non-absolute orthogonality of the evolved variant of *Methanocaldococcus jannaschii* tyrosyl-tRNA in *E. coli*.^30^

**Fig. 3.**
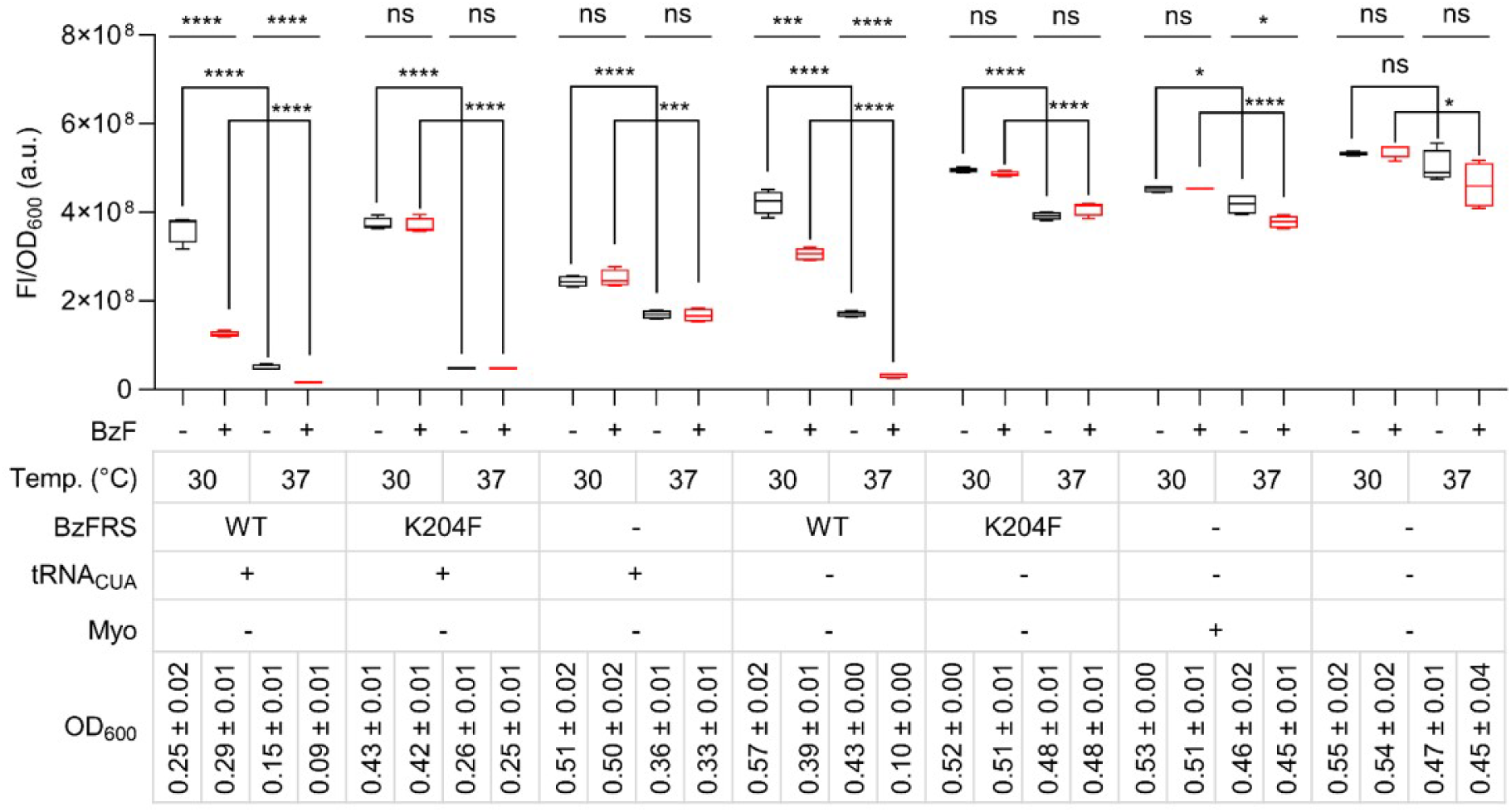
Impact of exogenous BzFRS and tRNA_CUA_ on temperature sensitivity in gene expression in DH10β Δ*cyaA*. sfGFP expression levels were evaluated by incubating DH10β Δ*cyaA*/F1RP-GFP/F2P cells, which contained one of the following plasmids: SP, SP-K204F, SP-aaRS^null^, SP-tRNA^null^, SP-K204F-tRNA^null^, SP^null^-Myo, or SP^null^. These are incubated in LB broth, both with and without BzF (0.5 mM) for 24 h at specified temperatures. WT refers to BzFRS and Myo represents sperm whale myoglobin. Blank plasmids were constructed by deleting either BzFRS (SP-aaRS^null^), tRNA_CUA_ (SP-tRNA^null^), or both (SP^null^). Additionally, SP-K204F-tRNA^null^ lacks tRNA_CUA_, derived from SP-K204F; n = 4; Box limits indicate the interquartile range; whiskers represent the range from minimum to maximum; the center line denotes the median; BzF conditions were highlighted in red; error bars stand for standard deviation; ns (not significant), * P < 0.05, *** P < 0.001, **** P < 0.0001 by Student’s t-test; and variability in OD_600_ values is represented by the standard deviation.

To investigate whether reduced catalytic activity of BzFRS influences temperature sensitivity, we engineered a down mutant, BzFRS-K204F, by mutating a single amino acid in the ATP-binding KMSSS loop. This modification resulted in a 104-fold decrease in amber suppression efficiency compared to wild-type BzFRS, as determined by the expression of the sfGFP-Y151TAG gene (Supplementary Fig. 5).^31^ As expected, the expression of BzFRS-K204F alone did not significantly affect temperature sensitivity, with only a 1.3-fold deviation observed between 30 °C and 37 °C in the absence of BzF, and a 1.2-fold deviation in its presence. This marginal effect might be due to the increased energy demand associated with the overexpression of exogenous genes. We observed a similar deviation when expressing sperm whale myoglobin (Myo) in DH10β Δ*cyaA*/F1RP-GFP/F2P cells under the control of the same *tac*I promoter^32^ (Fig. 3). This suggests that the overexpression of exogenous genes contributes to the temperature sensitivity of GOI expression to some extent.

Surprisingly, co-expression of BzFRS-K204F with tRNA_CUA_ led to significant temperature sensitivity, demonstrating a 7.7-fold deviation between 30 °C and 37 °C. This deviation is notably larger than that observed in the tRNA_CUA_-only condition, which was 2.5-fold. Considering the diminished ATP binding ability of the K204F variant, ATP consumption by BzFRS-K204F is likely not a contributing factor, further supported by the observation that the BzFRS-K204F/tRNA_CUA_ condition, similar to the tRNA_CUA_-only and BzFRS-K204F-only conditions, was unaffected by BzF. Therefore, we propose that BzFRS-K204F still can interact with tRNA_CUA_, inhibiting its function and resulting in a 1.5-fold increase in sfGFP fluorescence under the BzFRS-K204F/tRNA_CUA_ condition compared to the tRNA_CUA_-only condition at 30 °C. At 37 °C, BzFRS-K204F may lose its binding affinity towards tRNA_CUA_, ceasing to act as a tRNA_CUA_ scavenger. The additive effect of tRNA_CUA_ cross-reaction, BzFRS-K204F overexpression, and the marginal enzymatic activity of BzFRS-K204F collectively led to a 3.5-fold reduction in sfGFP fluorescence compared to the tRNA_CUA_-only condition (Fig. 3).

### Gene expression is temperature sensitive only for cAMP-dependent promoters in ECM-BACTH

In the subsequent phase of our study, we sought to understand the underlying cause of the temperature sensitivity observed in the presence of BzFRS/tRNA_CUA_. We postulated that the lower sfGFP expression at higher temperatures might be linked to a depletion of intracellular ATP levels, a scenario that could arise from ATP consumption by overexpressed aaRS and tRNA. To test this hypothesis, we performed experiments to correlate the sfGFP expression and intracellular ATP levels under different conditions (Fig. 4).

**Fig. 4.**
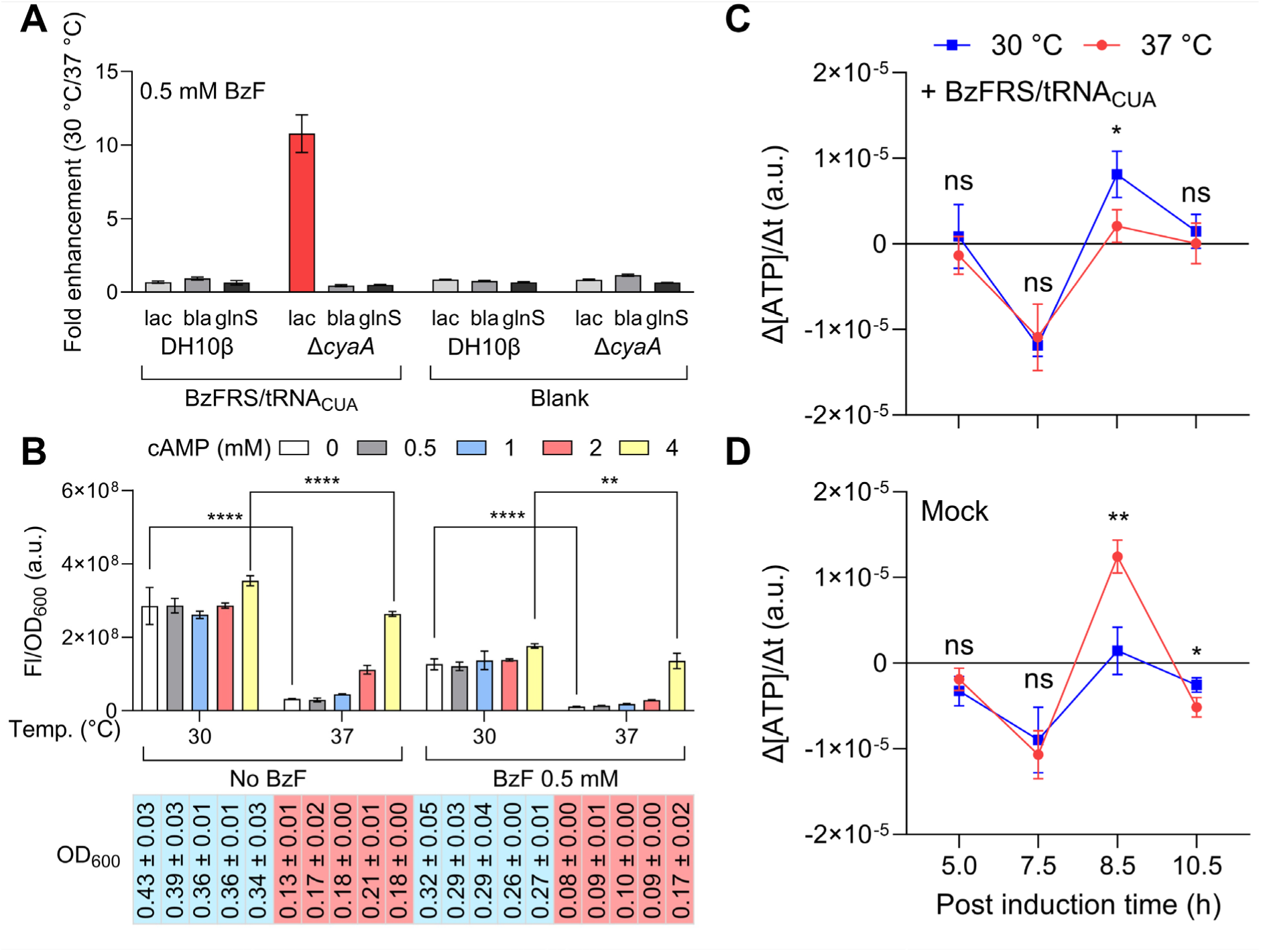
Correlation of the reduced gene expression with ATP and cAMP depletion. **A** Comparison of low-temperature-dependent gene expression levels controlled by three different promoters (*lac*, *bla*, and *glnS*), with or without BzFRS/tRNA_CUA_, in both DH10β and DH10β Δ*cyaA* strains. The data presented is a ratio of fluorescence intensities at 30 °C and 37 °C. **B** sfGFP fluorescence was measured after incubating DH10β Δ*cyaA*/F1RP-GFP/F2P/SP cells under various cAMP conditions (0, 0.5, 1, 2, and 4 mM) at 30 °C and 37 °C. Cells were incubated in LB broth both in the absence and presence of BzF (0.5 mM) for 24 h. **C**, **D** Comparative analysis of intracellular ATP level changes in DH10β Δ*cyaA*/F1RP-GFP/F2P cells carrying either the SP (**C**) or SP^null^ (**D**) plasmid. Cells were cultured in LB broth with BzF (0.5 mM) for 12 h. Conditions were set at 30 °C (represented in blue) or 37 °C (represented in red). At specified time points (3 h, 7 h, 8 h, 9 h, and 12 h), cells were collected; followed by cell lysis for ATP level determination via a luciferase-based ATP assay. Changes in ATP concentration slopes over time (Δ[ATP]/Δt) at specific time points are plotted. n = 4; ns (not significant), * P < 0.05, ** P < 0.01, **** P < 0.0001 by Student’s t-test; error bars, standard deviation; and variability in OD_600_ values is represented by the standard deviation.

To begin with, we evaluated the gene expression levels governed by three distinct promoters (*lac*, *bla*, and *glnS*) in the context of either the presence or absence of BzFRS/tRNA_CUA_. These investigations were carried out using both DH10β and DH10β Δ*cyaA* strains, with temperature settings at 30 °C and 37 °C. Of note, the *lac* promoter, which is known to be cAMP-dependent,^22^ exhibited the most significant temperature-dependent change. These changes represented a 9-fold and 11-fold enhancement in expression at the lower temperature (30 °C) relative to that at 37 °C in the DH10β Δ*cyaA* strain expressing BzFRS/tRNA_CUA_ in the absence or presence of BzF, respectively (Fig. 4A and Supplementary Figs. 6–8).

Subsequently, we conducted experiments with DH10β Δ*cyaA*/F1RP-GFP/F2P/SP cells at both 30 °C and 37 °C, spanning a range of cAMP conditions, with and without BzF. Remarkably, we observed that the diminished sfGFP expression at the higher temperature (37 °C) could be rescued by the addition of cAMP, a pattern evident regardless of the presence of BzF. When we supplemented with a high concentration of cAMP (4 mM), the reduced sfGFP expressions at 37 °C were restored up to 75% or 77% in the absence or presence of BzF, respectively (Fig. 4B). However, it is important to note that the expression levels under the control of the other two promoters (*bla* and *glnS*), which are not cAMP-dependent,^33^ remained unaffected by the addition of cAMP (Supplementary Fig. 9). These findings provide additional support for our hypothesis that the observed temperature sensitivity was primarily due to lower cAMP levels at higher temperatures.

### Expression of BzFRS/tRNA_CUA_ elevates ATP consumption rates at higher temperatures in ECM-BACTH

We evaluated the intracellular ATP levels and sfGFP expression in DH10β Δ*cyaA*/F1RP-GFP/F2P cells harboring either the SP or SP^null^ plasmid. After incubating these cells in LB broth, with or without BzF, for 12 h, they were subjected to analysis for sfGFP fluorescence and intracellular ATP levels (Supplementary Fig. 10). When cultured with BzF, the rate of intracellular ATP concentration changes over time (Δ[ATP]/Δt) remained similar between two temperatures (30 °C and 37 °C) until 8 hours post-induction (hpi). However, a significant deviation in the rate of ATP concentration change was observed around 8.5 hpi. As illustrated in Fig. 4C and 4D, it became evident that Δ[ATP]/Δt showed lower values at 37 °C compared to 30 °C in the presence of BzFRS/tRNA_CUA_, contrasting with the control. In the absence of BzF, Δ[ATP]/Δt consistently has lower values at 37 °C compared to 30 °C, but this effect was observed exclusively in the presence of BzFRS/tRNA_CUA_ expression, as illustrated in Supplementary Figs. 10 and 11. We attribute the consistently more negative values in Δ[ATP]/Δt at 37 °C compared to 30 °C, observed around 8.5 hpi, to a higher rate of ATP consumption relative to ATP biosynthesis in the presence of BzFRS/tRNA_CUA_ at elevated temperatures. This supports our hypothesis that the temperature sensitivity induced by exogenous aaRS/tRNA is related to elevated ATP consumption rate at higher temperature. Moreover, it is worth noting that these conditions could also impact ATP biosynthesis, given that the gene is regulated by cAMP–CRP.^22^

In further observations, the suppressive effect of BzFRS/tRNA_CUA_ on sfGFP expression at 37 °C was less prominent without BzF until 12 h, as seen in Supplementary Figs. 10A and 12B. However, when BzF was present, the suppression became more efficient, as highlighted in Supplementary Figs. 10C and 12D. Notably, the detrimental effect of BzF on sfGFP expression was visible only in the presence of BzFRS. When BzF was present, sfGFP expression levels were generally reduced in the presence of BzFRS alone at both temperatures (1.4-fold at 30 °C and 5.3-fold at 37 °C). A similar pattern of reduction was observed with the BzFRS/tRNA_CUA_ pair (2.9-fold at 30 °C and 3.0-fold at 37 °C). However, conditions with tRNA_CUA_ alone remained unaffected by BzF (Fig. 3). This suggests that the aminoacylation reaction mediated by BzFRS in the presence of BzF, potentially leading to ATP depletion, significantly contributes to the reduction in sfGFP expression.

The increased ATP consumption rate during the early growth phase significantly affects cell growth. Specifically, DH10β Δ*cyaA*/F1RP-GFP/F2P/SP cells exhibited markedly slower growth, with a 6.6-fold (without BzF) and 8.3-fold (with BzF) reduction in OD_600_ after a 12-hour culture at 37 °C, compared to their SP^null^ counterparts (Supplementary Fig. 12A and 12C). Moreover, while the cells with the ECM-BACTH system expressing SP showed slower growth at 37 °C than at 30 °C, those expressing SP^null^ exhibited the inverse trend (Supplementary Fig. 12A and 12C). This discrepancy could stem from the suppressed expression of approximately 380 cAMP-dependent genes under the additional ATP-consuming conditions induced by BzFRS/tRNA_CUA_, some of which could be pivotal for growth.^22, 34, 35^ Notably, the overexpression of BzFRS-K204F and Myo did not adversely affect cell growth at either temperature (30 °C or 37 °C), suggesting that the mere overexpression of heterologous genes is not the primary cause of growth defects (Fig. 3 and Supplementary Table 3). Intriguingly, despite observing a temporary increase in the ATP consumption rate at a specific time point (around 8.5 hpi) at 37 °C, a consistent finding in our study was the increased level of intracellular ATP in cells overexpressing BzFRS/tRNA_CUA_ compared to the control (Supplementary Fig. 10). The accumulation of ATP suggests inefficiencies in ATP utilization by essential cellular functions in the presence of the exogenous aaRS/tRNA, contributing to the observed growth defects.

### Mechanistic insights into ATP consumption in genetic code expanded system: a consideration for hosts with compromised ATP homeostasis

Based on our findings, we propose a couple of mechanistic insights leading to increased ATP consumption, contributing to the temperature sensitivity observed in our system. Firstly, although the effect may be marginal, the overexpression of exogenous genes, such as orthogonal aaRS (aaRS^O^), could contribute to ATP depletion at higher temperatures through the general aminoacylation process. Secondly, the simultaneous overexpression of aaRS^O^ and orthogonal tRNA (tRNA^O^), encoded on the suppressor plasmid, might exacerbate ATP drain, especially at elevated temperatures where metabolic activities are intensified. Lastly, the imperfect orthogonality of the introduced aaRS^O^/tRNA^O^ system could lead to unintended cross-reactions with native *E. coli* aaRS/tRNAs or with canonical amino acids, triggering additional ATP-dependent aminoacylation reactions and further depleting ATP reserves (Supplementary Fig. 13). In addition to the cross-reactions, other potential mechanisms, such as non-productive aminoacylation or the production of diadenosine tetraphosphate without tRNA interaction, should be considered when aaRS^O^ is expressed alone.^19^

These mechanisms cumulatively exert a significant metabolic burden, resulting in ATP and cAMP shortage at higher temperatures, which eventually impacts the expression of the GOI. This highlights a critical aspect of implementing GCE systems, particularly in hosts with compromised ATP homeostasis, such as the *cyaA* knockout system. Interestingly, in the conventional protein expression strain BL21(DE3), a comparable effect was observed, but it was specifically induced by the presence of BzF alongside the BzFRS/tRNA_CUA_ pair, resulting in a 2.8-fold and 36.9-fold decrease in sfGFP isolated yield at 30 °C and 37 °C, respectively, when compared to the control (Supplementary Fig. 14, see supporting information for details). This aligns with previous report that modified strains like BL21 are vulnerable to orthogonal translation systems.^18^ Therefore, these findings underscore the importance of careful system design and component selection, with an emphasis on perfect orthogonality, to minimize the metabolic load and ensure robust functioning of the system.

### Leveraging temperature sensitivity in a dual-input biologic gate

We further expanded on our observation of temperature sensitivity by developing a dual-input biologic gate. This setup uses the BzF and temperature as binary inputs (Fig. 5A). We constructed this system by replacing ECM with its BzF-dependent variant (ECB) and adding a suppressor plasmid SP, thereby establishing the ECB-BACTH (Supplementary Fig. 15). Then, we asked whether this genetic circuit could operate as an AND gate with gene expression serving as the output. To evaluate this, we cultured DH10β Δ*cyaA*/F1RP*-CAT/F2P*/SP and DH10β Δ*cyaA*/F1RP*-GFP/F2P*/SP cells on LB-agar under specified conditions for 72 h (* denotes replacing ECM with ECB). Following this, we examined the growth of colonies in the presence of Cm and the fluorescence of sfGFP under conditions represented by the binary codes 00, 01, 10, and 11. Notably, in this setup, the GOIs are only expressed when both conditions are met: in the presence of BzF (X_1_ = 1) and at a lower temperature (30 °C; X_2_ = 1) (Fig. 5B, 5C, and Supplementary Figs. 16–18). Further, in liquid culture, sfGFP fluorescence under the permissive condition (binary code = 11) was markedly higher than other conditions, showing a 36-fold enhancement compared to the 01 condition (Fig. 5D). Interestingly, fluorescence under the 01 condition was lower than under both the 00 and 10 conditions, each of which showed about a 2-fold increase compared to the 01 condition. However, it’s important to note that the signals in the 00 and 10 conditions are still significantly lower than that of the 11 condition (Fig. 5D). This diminished fluorescence in the 01 condition is likely due to reduced leaky expression at lower temperatures.

**Fig. 5.**
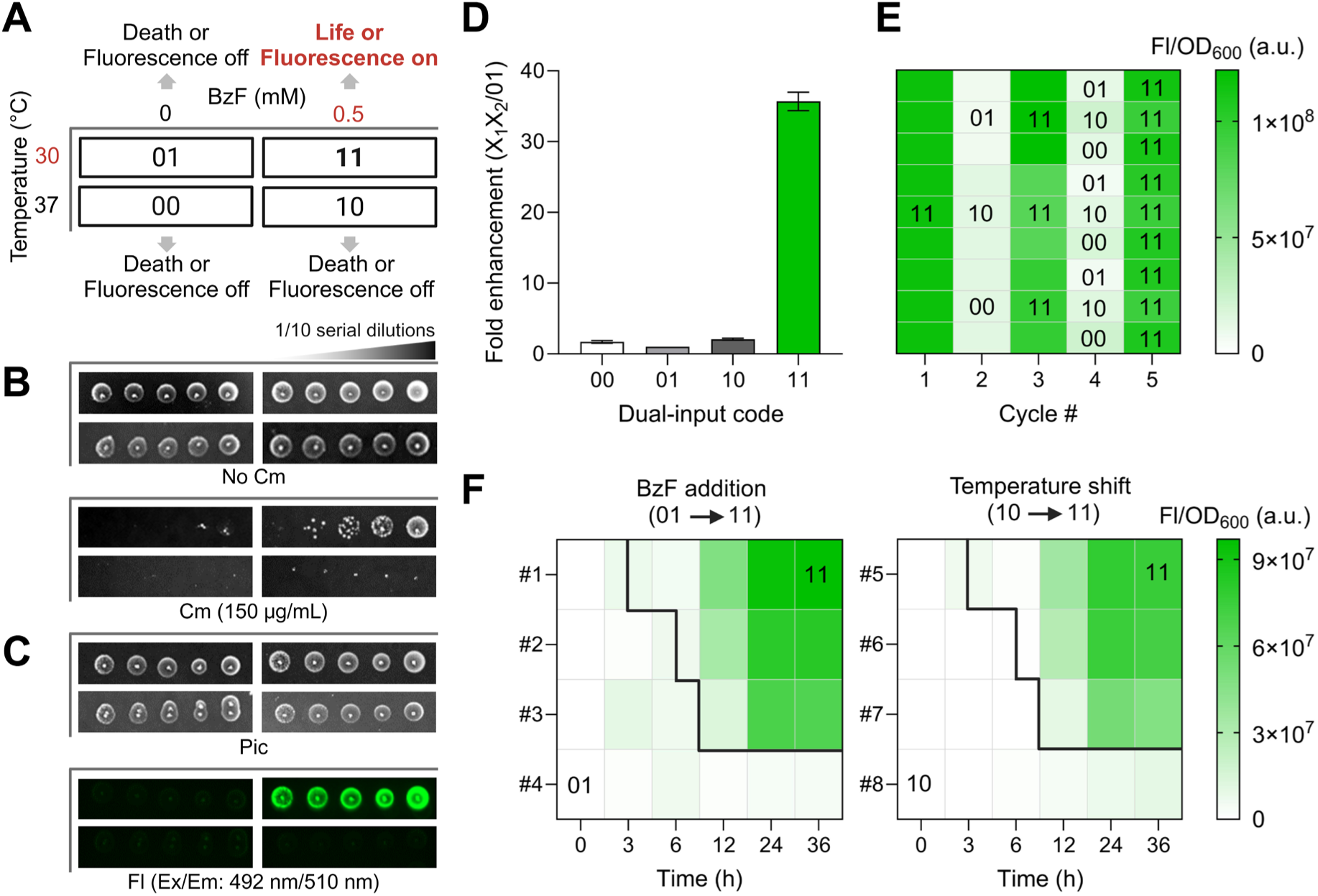
A dual-input biologic gate. **A** Schematic overview of a dual-input biologic gate utilizing presence (X_1_ = 1) or absence (X_1_ = 0) of BzF (0.5 mM) and low (30 °C, X_2_ = 1) or high (37 °C, X_2_ = 0) temperature as an AND gate binary code. The GOI is expressed only if both conditions (presence of BzF and 30 °C) are met. All the conditions with the binary codes (00, 01, 10, and 11) are shown in the box. **B** Dilution spot images of the DH10β Δ*cyaA*/F1RP*-CAT/F2P*/SP cells on LB-agar, both in the absence and presence of Cm (150 μg/mL), incubated under specified conditions for 72 h. **C** Dilution spot images (Pic) of the DH10β Δ*cyaA*/F1RP*-GFP/F2P*/SP cells on LB-agar incubated under specified conditions for 72 h. The sfGFP fluorescence (Fl) was measured at the same time. **D** DH10β Δ*cyaA*/F1RP*-GFP/F2P*/SP cells were incubated for 24 h under four conditions (00, 01, 10, and 11). sfGFP fluorescence intensities, normalized to the non-permissive condition (01), are shown. Error bars denote standard deviation (n = 4). **E** DH10β Δ*cyaA*/F1RP*-GFP/F2P*/SP cells were incubated for 24 h under permissive conditions (0.5 mM BzF and 30 °C). Then, the washed culture was diluted 100-fold to fresh LB broth and grown under three different non-permissive conditions (00, 01, and 10) for 24 h. Additional cultures were examined by alternating between permissive (11) and non-permissive conditions (00, 01, and 10). The level of GOI expression was assessed by measuring sfGFP fluorescence. **F** DH10β Δ*cyaA*/F1RP*-GFP/F2P*/SP cells cultured for 24 h under non-permissive conditions (00) were diluted 200-fold into non-permissive media (01, left heat map or 10, right heat map). Subsequently, the sfGFP expression was induced by shifting to permissive conditions (11), either by the addition of BzF (0.5 mM) or lowering the temperature (from 37 °C to 30 °C). This shift was conducted at 3 hours (#1 and #5), 6 hours (#2 and #6), or 9 hours (#3 and #7) post-inoculation. Samples that did not undergo these shifts served as controls (#4 and #8). Two different culture conditions (01 and 11; 10 and 11) are separated by black lines. The presented data is an average of biological replicates (n = 4).

The sfGFP expression level was typically increased as the temperature decreased. In fact, at 25 °C with 0.5 mM BzF (input code 12), we observed an 86-fold enhancement in normalized sfGFP fluorescence compared to that with input code 02 (Supplementary Fig. 1). This enhancement might be attributed to further reduced ATP consumption at this temperature. However, despite the pronounced dynamic range observed at 25 °C, we selected 30 °C as the standard permissive temperature. This decision was based on the observation that growth under the condition denoted by code 11 (30°C) had the highest growth rate (Supplementary Fig. 19B), with a doubling time of 88 ± 10 min (Supplementary Table 3). Collectively, these results affirm that the novel biologic gate effectively modulates gene expression based on a specific set of conditions, offering an additional layer of control in genetic engineering applications.

### Switchability of the dual-input biologic gate: towards precise control of gene expression

To assess the switchability of our dual-input biologic gate, toggling GOI expression on or off by altering conditions, we incubated DH10β Δ*cyaA*/F1RP*-GFP/F2P*/SP cells under permissive conditions (presence of BzF at 30 °C, binary code = 11) for 24 h. Afterward, the culture was washed and transferred to three different non-permissive environments (binary codes = 00, 01, and 10). Finally, we could effectively modulate the expression of the GOI by alternating between these permissive and non-permissive conditions (Fig. 5E).

We further showcased this switchability by initially placing the cells to non-permissive conditions, specifically in the absence of BzF at 30 °C (binary code = 01) or in the presence of BzF at 37 °C (binary code = 10). Upon modifying the environmental conditions to permissive ones, either by introducing BzF (0.5 mM) or lowering the temperature to 30 °C, the sfGFP expression was induced, signifying an on-state of the genetic switch (Fig. 5F). This control of gene expression, driven by environmental cues, opens avenues for potential advancements in synthetic biology applications where orthogonal modules for gene expression is needed.

### Three-input AND gate using ncAA, temperature, and small molecule inducer

Utilizing our circuit design that relies on the IPTG induction for both the target gene and BzFRS expression, we tested whether the IPTG acts as an additional input (Fig. 6A). As a result, we confirmed that the additional requirement confers tighter regulation of the target gene expression. In fluorescence measurements after culturing DH10β Δ*cyaA*/F1RP*-GFP/F2P*/SP cells on LB for 24 h, we observed 42-fold enhancement in sfGFP fluorescence intensity under the permissive condition (input code = 111) compared to the non-permissive condition (input code = 110). The leakiest non-permissive condition was 001, exhibiting a 3-fold higher fluorescence compared to the 110 condition (Fig. 6B). This could be attributed to the inherent leakiness of the *lac* promoter in the presence of IPTG, which might be accelerated at higher temperatures (e.g., 37 °C) in the absence of metabolic burden through BzF. Nonetheless, all non-permissive conditions showed significantly lower sfGFP expression levels compared to the permissive condition (input code = 111).

**Fig. 6.**
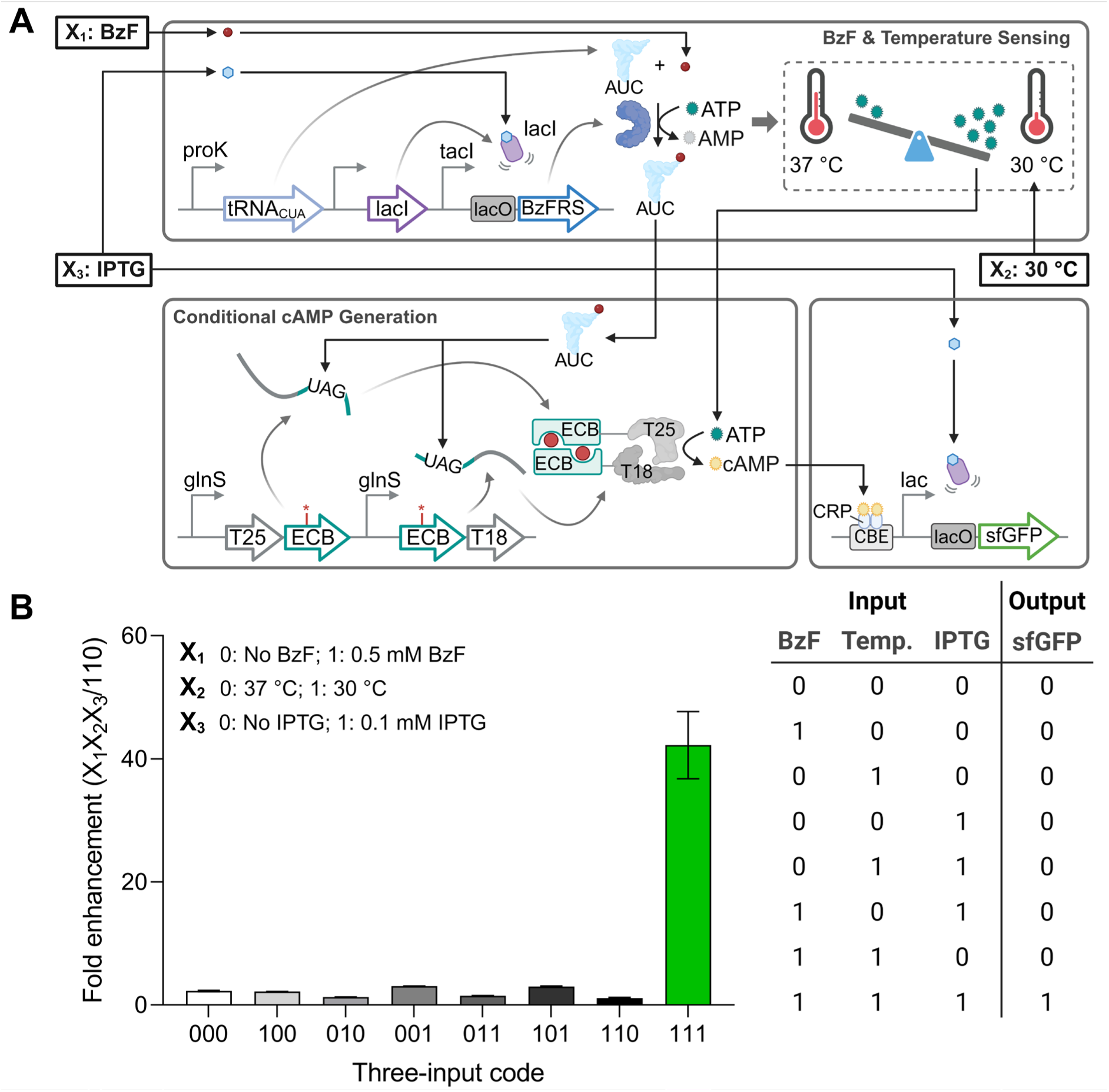
A three-input biologic gate. **A** Schematics of the genetic circuit flow that generates sfGFP fluorescence in response to the three binary inputs (X_1_X_2_X_3_). X_1_ represents the absence (0) or presence (1) of 0.5 mM BzF; X_2_ represents the temperature conditions, with 0 indicating 37 °C and 1 indicating 30 °C; and X_3_ represents the absence (0) or presence (1) of 0.1 mM IPTG. **B** The fold enhancement of sfGFP expression is presented as a bar graph. DH10β Δ*cyaA*/F1RP*-GFP/F2P*/SP cells were incubated for 24 h under eight different conditions (000, 100, 010, 001, 011, 101, 110, and 111). Pronounced sfGFP expression (output code = 1) was observed exclusively when input code 111 was applied. The sfGFP fluorescence intensities were normalized to the non-permissive condition (110). Error bars denote standard deviation (n = 4).

## Discussion

The development of biological logic gates for precise control of gene expression remains an ongoing challenge. Various synthetic modules for circuit design have been explored,^28^ including the use of amber suppressors,^36^ orthogonal ribosome/orthogonal mRNA pairs,^37^ genetic code expansion,^38, 39^ and small RNAs.^40^ Achieving this control relies on the development of genetic circuits that can effectively respond to externally applicable stimuli while maintaining compatibility with one another. Temperature serves as a valuable input signal due to its non-invasive nature and reversibility.^41^ However, the commonly used temperature-sensitive variant of the bacteriophage λ repressor, cI857, comes with a notable drawback. Its heat-inducible nature may not align with the physiological conditions of bacteria.^42^ Consequently, researchers have been exploring alternative cold-inducible switch systems,^41, 43, 44^ although they remain notably rare.

We have developed an AND gate that utilizes a novel temperature- and ncAA-controlled gene expression system. Our system offers several advantages compared to conventional methods used in AND gates. First, we have introduced a novel ncAA responsive module into the biological logic gate, expanding the scope of external stimuli, in addition to the rare examples.^38, 39^ Second, we showcased a new cold-inducible system, addressing potential issues with the stress effects and physiological responses associated with heat shock.^42^ Lastly, unlike previous systems vulnerable to mutations (e.g., strategies with simple amber codon insertion), our system remains robust due to its absolute dependency on BzF for dimer formation.^25^ Overall, our system will provide additional toolkits in conjunction with other biologic gates.

In conclusion, the serendipitous discovery of temperature sensitivity in cAMP-dependent gene expression led us to innovatively design a novel biological AND gate, leveraging the GCE strategy. The insights we gleaned from the associated metabolic burden of overexpressing exogenous aaRS and tRNA, and their effect on ATP consumption, have underlined the importance of careful system design and component selection. From this perspective, acknowledging the current limitations of our biological logic gate, particularly its broader impact on cellular processes, is crucial for our ongoing research. Despite these challenges, the orthogonal nature of ECB presents a promising pathway. In light of this, our future efforts will focus on using the ECB module to induce the reassembly of split proteins, other than adenylate cyclase, thus avoiding the production of metabolites such as cAMP. This aims to minimize broader cellular impacts and create more universally applicable biological devices.

## Methods

### Reagents

The restriction enzymes NdeI, PstI, KpnI, and T4 DNA ligase were purchased from Enzynomics (South Korea), while DpnI was obtained from Toyobo (Japan). Primers were acquired from Macrogen (South Korea), and for PCR experiments, we used TOPsimple™ PCR PreMIX-Forte from Enzynomics. Site-directed mutagenesis was facilitated using both the EZchange™ Site-directed Mutagenesis kit (Enzynomics) and the KOD-Plus Mutagenesis Kit (Toyobo). PCR products and products from restriction digestion were purified by agarose gel electrophoresis, the LaboPass™ Gel Extraction Kit (Cosmogenetech, South Korea), the Gel DNA Clean-up Kit, the PCR DNA Clean-up Kit (both from SmartGene, South Korea), and the DNA Clean & Concentrator (Zymo Research, USA). Plasmid purifications were executed with the EZ-Pure™ Plasmid Prep Kit Ver. 2 from Enzynomics and the Plasmid Mini Prep Kit from SmartGene. DNA Sanger sequencing was performed by Macrogen. Ampicillin, spectinomycin, tetracycline, chloramphenicol, and bovine serum albumin were purchased from MilliporeSigma (USA). Kanamycin, L-(+)-arabinose, and adenosine 3’, 5’-cyclic monophosphate (cAMP) were purchased from Tokyo chemical industry (TCI, Japan). β-D-1-thiogalactopyranoside (IPTG), sodium hydroxide, sodium chloride, sodium phosphate monobasic anhydrous, sodium phosphate dibasic anhydrous, and magnesium sulfate heptahydrate were purchased from Duksan (South Korea). Ethyl alcohol, methyl alcohol, and imidazole were purchased from Daejung (South Korea). *p*-benzoylphenylalanine (BzF) was purchased from Combi-Blocks, Inc. (USA). BacTiter-Glo™ Microbial Cell Viability Assay reagent was purchased from Promega (USA). We purchased B-PER™ Complete Bacterial Protein Extraction Reagent and Pierce™ 660 nm Protein Assay Reagent from Thermo Fisher Scientific (USA), Ni-NTA Agarose from QIAGEN (Germany), Ni-NTA Magnetic Beads from Bioneer (South Korea) and Amicon Ultra Centrifugal Filter Units from MilliporeSigma. Notably, the Gibson assembly master mix was custom-prepared using components including 1 M Tris-HCl (Enzynomics), magnesium chloride (Daejung), 100 mM dNTPs (Enzynomics), DL-dithiothreitol (TCI), polyethylene glycol 8,000 (MilliporeSigma), β-nicotinamide adenine dinucleotide sodium salt (MilliporeSigma), T5 exonuclease (Enzynomics), Phusion HF DNA polymerase (New England Biolabs, NEB, USA), and Taq ligase (Enzynomics).

### E. coli strain

Plasmids and primers utilized in this study can be found in Supplementary Tables 1 and 4, respectively. The *E. coli* strains utilized in this study included DH10β (NEB10β), sourced from NEB, and BL21(DE3), purchased from Enzynomics. In this research, the DH10β Δ*cyaA* strain was established via λ-Red mediated recombineering using the pKD46^45^ plasmid. The *cyaA* gene deletion was based on the Keio Collection, specifically deleting residues 2-842 of the *cyaA* gene. The *cyaA* gene was substituted with a tetracycline resistance gene, which was amplified from pKD4-tet^R^ with primers oMK-56 and oMK-57. The successful deletion was verified through colony PCR, and tetracycline (12.5 μg/mL) was used for strain selection (Supplementary Fig. 20).

## Plasmid construction

### F1RP-CAT and F1RP*-CAT

The ECM gene was amplified from the plasmid pKTECM^25^ using primers oMK-16/oMK-19 and oMK-17/oMK-18 to give fragment A and B, respectively. Likewise, the ECB gene was amplified from the plasmid pKTECB^25^ using primers oMK-16/oMK-19 and oMK-17/oMK-112 to give fragment A* and B*, respectively. The fragments A and A* were digested with PstI and then ligated to the PstI fragment of pT25SC-rep, which originated from p25N,^46^ resulting in the creation of F1RP-CAT and F1RP*-CAT, respectively.

### F2P and F2P*

Fragments B and B* were digested using NdeI and KpnI and subsequently ligated to the NdeI and KpnI fragment of pBK-SCT18, which was sourced from pUT18,^46^ resulting in F2P and F2P*, respectively.

### F1RP^null^-CAT

F1RP^null^-CAT was generated via site-directed mutagenesis PCR (utilizing oMK-110/oMK-111).

### F1RP-GFP, F1RP*-GFP, and F1RP^null^-GFP

Gibson assembly was used to replace the reporter gene from the chloramphenicol acetyltransferase (CAT) gene to the super-folder green fluorescence protein (sfGFP) gene. The gene for sfGFP was amplified from the plasmid pET22b-T5-sfGFP^47^ using primers oMK-132 and oMK-133. To construct F1RP-GFP, F1RP*-GFP, and F1RP^null^-GFP, the backbones of F1RP-CAT, F1RP*-CAT, and F1RP^null^-CAT were amplified with oMK-134 and oMK-135. Then, sfGFP fragment and backbones fragment were ligated by Gibson assembly to give F1RP-GFP, F1RP*-GFP, and F1RP^null^-GFP.

### F1RP-GFP-bla and F1RP-GFP-glnS

The *bla* promoter was amplified from the F1RP-GFP plasmid using primers oMK-227 and oMK-228, while the sfGFP gene was obtained with primers oMK-133 and oMK-225. These fragments were combined through overlap extension PCR to generate the bla-sfGFP. To construct F1RP-GFP-bla, the bla-sfGFP fragment and a PCR product from F1RP-CAT (amplified using oMK-226 and oMK-160) were ligated via Gibson assembly. To construct the glnS version, the sfGFP PCR fragment from the pET22b-T5-sfGFP plasmid was amplified using primers oMK-213 and oMK-214. After digestion with NdeI and PstI, it was inserted into the NdeI/PstI site of pBK-SCT18, yielding the glnS-sfGFP construct. The glnS-sfGFP gene was then amplified using oMK-133 and oMK-223, while the backbone of F1RP-CAT was obtained with primers oMK-160 and oMK-222. The glnS-sfGFP fragment and the F1RP-CAT backbone were combined via Gibson assembly to produce the F1RP-GFP-glnS.

### SP-tRNA^null^, SP-aaRS^null^, SP^null^, and SP^null^-Myo

SP-tRNA^null^, SP-aaRS^null^, SP^null^, and SP^null^-Myo were constructed based on the SP (pUltra-BzF) plasmid.^25^ These constructs were generated through site-directed mutagenesis PCR employing primers oMK-180/oMK-200 for SP-tRNA^null^, oMK-181/oMK-186 for SP-aaRS^null^, and oMK-180/oMK-181 for SP^null^. The backbone of SP^null^-Myo was amplified from the plasmid SP-tRNA^null^ with oMK-465 and oMK-466. The gene for sperm whale myoglobin was amplified from the plasmid pET22b-T5-Myo^48^ using primers oMK-467 and oMK-468. Then, backbone fragment and myoglobin fragment were ligated by Gibson assembly to give SP^null^-Myo.

### SP-K204F and SP-tRNA^null^-K204F

SP-K204F and SP-K204F-tRNA^null^ were generated via site-directed mutagenesis PCR from the SP and SP-tRNA^null^, respectively, using primers oMK-0530/oMK-0531.

### Culture condition

Antibiotics were used at the following concentrations unless otherwise stated: ampicillin (Amp, 100 μg/mL), kanamycin (Kan, 50 μg/mL), spectinomycin (Spec, 50 μg/mL), and tetracycline (Tet, 12.5 μg/mL). For the reporter gene CAT, a range of chloramphenicol (Cm) concentrations, including 50, 100, 150, and 200 μg/mL, were tested.

The optimal concentrations of IPTG (0.1 mM) and BzF (0.5 mM) were determined by assessing the growth of the DH10β Δ*cyaA* containing F1RP*-CAT/F2P*/SP in LB supplemented with Amp, Kan, Spec, and Cm (50 μg/mL), under different conditions involving combinations of 0.5 mM and 1 mM BzF and 0.1 mM and 1 mM IPTG (Supplementary Fig. 21A). The optimal BzF concentration was further determined through fluorescence measurements using the DH10β Δ*cyaA* harboring F1RP*-GFP/F2P*/SP. A series of half dilutions, starting from a BzF concentration of 1 mM, was carried out to determine the concentration producing the highest fluorescence intensity (Supplementary Fig. 21B). To assess the toxicity of the BzF, OD_600_ was measured for DH10β Δ*cyaA*/F1RP-CAT/F2P cells harboring either the SP or SP^null^ plasmid, incubated in LB broth at various BzF concentrations (0, 0.1, 0.5, and 1 mM) and temperatures (30 °C and 37 °C) over 24 h. Bacterial growth remains unaffected by BzF at concentrations as high as 1 mM (Supplementary Fig. 22).

Seed cultures were incubated in LB medium at 37 °C for 24 h, shaking at 180 rpm. Subsequently, the seed cultures were diluted 200-fold for liquid cultures, and for solid cultures, a dilution series of transformants (approximately 10^6^, 10^5^, 10^4^, 10^3^, and 10^2^ cells) was prepared and spotted onto LB-agar plates. Transformants without SP, SP-tRNA^null^, SP-aaRS^null^, SP^null^, SP-K204F, SP-K204F-tRNA^null^, or SP^null^-Myo plasmids were grown in LB or LB-agar medium supplemented with Amp and Kan. For cultivating transformants containing SP or its variant plasmids, additional supplements of Spec and IPTG were included.

### Growth experiment

Growth experiments for assessing the Cm resistance were conducted either by culturing DH10β Δ*cyaA* transformants on solid LB-agar or liquid LB. For the transformants harboring F1RP-CAT/F2P, F1RP^null^-CAT/F2P, F1RP-CAT/F2P^null^, or F1RP^null^-CAT/F2P^null^, we tested range of Cm concentrations (0, 50, 100, and 200 μg/mL) at 37 °C for 20 h. Transformants containing F1RP-CAT/F2P/SP, F1RP-CAT/F2P/SP^null^, F1RP*-CAT/F2P*/SP were incubated for 72 h on LB-agar plates with Cm (0, 100, 150, or 150 μg/mL) and BzF (0 or 0.5 mM) at two different temperatures (30 °C and 37 °C). The dilution spot images were taken with ChemiDoc™ MP (Bio-Rad, USA). Liquid cultures of DH10β Δ*cyaA* carrying the F1RP*-CAT/F2P*/SP were subjected to conditions similar to the solid cultures, beginning with a 200-fold dilution from the 24 h seed cultures. Optical densities at 600 nm (OD_600_) were measured using either an OD_600_ DiluPhotometer™ (Implen, Germany) or a SpectraMax iD3 Multi-Mode Microplate Reader (Molecular Devices, USA).

### Fluorescence assay

Fluorescence intensities of the bacterial strains were measured after incubating under the designated culture conditions. Both OD_600_ and fluorescence (Fl; excitation/emission: 485 nm/525 nm; exposure time: 140 ms) were measured using the SpectraMax iD3 Multi-Mode Microplate Reader. Subsequently, sfGFP fluorescence was normalized to cell count, represented as Fl/OD_600_. For temperature-based fluorescence intensity screening, the DH10β Δ*cyaA* strain carrying F1RP*-GFP/F2P*/SP was incubated across a temperature range (25 °C, 27 °C, 30 °C, 33 °C, 35 °C, and 37 °C), both with and without BzF (0.5 mM). Fluorescence images of *E. coli* on LB-agar plates were captured using ChemiDoc™ MP at excitation/emission: 492 nm/510 nm with an exposure time of 20 ms.

### cAMP titration

DH10β Δ*cyaA* cells equipped with the F1RP-GFP/F2P/SP plasmids, along with its variants F1RP-GFP-bla/F2P/SP and F1RP-GFP-glnS/F2P/SP, were cultured for 24 h. These strains were exposed to varying concentrations of cAMP (0, 0.5, 1, 2, and 4 mM), derived from a 1 M stock solution prepared in 2 M NaOH (aq.). The cells were grown in LB broth at both 30 °C and 37 °C for 24 h, with both the absence and presence of 0.5 mM of BzF. After the incubation period, the fluorescence intensities of the bacterial cultures were measured using the SpectraMax iD3 Multi-Mode Microplate Reader.

### Analysis of intracellular ATP contents

The intracellular ATP levels in *E. coli* were determined using the BacTiter-Glo Microbial Cell Viability Assay Reagent. A pre-treatment buffer was composed of 20 mM Tris-HCl (pH 7.4), 50 mM MgSO_4_, 4 mM EDTA, and 50% methanol.^49^ *E. coli* culture samples were prepared at intervals post-inoculation: 3, 7, 8, 9, and 12 h. These samples had their OD_600_ and fluorescence measured, and were then centrifuged at 4 °C at 3,000 g for 10 min. The supernatants were removed, and the obtained pellets were stored at −78 °C. For analysis, pellets were re-suspended in the pre-treatment buffer and subjected to a 30-min incubation at 70 °C in a heat block. Subsequent to this incubation, the solutions underwent centrifugation at 4 °C and 10,000 g for 5 min. Supernatants (25 μL) were combined with an equal volume of BacTiter-Glo Microbial Cell Viability Assay Reagent in a 384-well plate. This mixture was then agitated gently on an orbital shaker for 5 min at room temperature. Luminescence was recorded using the SpectraMax iD3 Multi-Mode Microplate Reader. The intracellular ATP content in *E. coli* was inferred from the ATP standard curve and then normalized by the OD_600_ value. For the ATP standard curve, a 5-fold dilution series ranging from 10 μM to 25.6 pM ATP in the pre-treated buffer was employed.

### Protein expression and mass spectrometry

Three distinct cultures of DH10β cells were conducted, each with a distinct plasmid set: pET22b-T5-sfGFP*^4^/SP, pET22b-T5-sfGFP*/SP-aaRS^null^, and pET22b-T5-sfGFP*. These cells were pre-cultured at 37 °C for 12 h, diluted 200-fold into LB broth supplemented with BzF (0 or 0.5 mM) and specific antibiotics at 37 °C. When the OD_600_ reached 0.4, IPTG (0.1 mM) was added to induce protein expression, and the cultures were further incubated at 30 °C for 16 h with shaking at 180 rpm.

To analyze the mass of the sfGFP variants, the cells were harvested by centrifugation at 4,000 g for 20 min. Cell pellets were resuspended in B-PER™ Complete Bacterial Protein Extraction Reagent and incubated on a rocking platform at room temperature for 15 min. The lysates were then centrifuged at 10,000 g for 20 min at 4 °C, and the resulting supernatants were loaded onto a Ni-NTA agarose column that had been pre-equilibrated with buffer A (100 mM NaH_2_PO_4_, pH 7.8, 100 mM NaCl, and 20 mM imidazole) and incubated for 1 h at 4 °C. After washing the column with buffer A and buffer B (100 mM NaH_2_PO_4_, pH 7.8, 100 mM NaCl, and 50 mM imidazole), the proteins were eluted using buffer C (100 mM NaH_2_PO_4_, pH 7.8, 100 mM NaCl, and 250 mM imidazole). The buffer was then exchanged to PBS (pH 7.4) using an Amicon Ultra 10 kDa centrifugal filter. The protein samples were analyzed using an Agilent Technologies 6530 accurate-mass quadrupole time-of-flight (Q-TOF) LC/MS instrument at the Core Research Facility of Pusan National University.

The ability for aminoacylation by BzFRS and BzFRS-K204F was indirectly compared by measuring sfGFP fluorescence in DH10β/pET22b-T5-sfGFP* cells harboring either SP or SP-K204F, incubated under identical conditions as described above.

To measure protein yield, we employed BL21(DE3)/F1RP-GFP/F2P strains, either containing SP or SP^null^. These strains were initially pre-cultured at 37 °C for 12 h, followed by a 200-fold dilution into LB broth enriched with specific antibiotics, and further incubated at 37 °C in the absence or presence of 0.5 mM BzF. After incubating for three hours, we added IPTG (0.1 mM) to initiate protein expression. The cultures were then incubated for an additional 5 h at either 30 °C or 37 °C, with shaking at 180 rpm. Separately, DH10β Δ*cyaA* cells were subjected to the same initial pre-culture and incubation steps, but IPTG (0.1 mM) was added at the inoculation step, and the incubation period was extended to 24 h. Protein purification from both sets of cultures utilized Ni-NTA magnetic beads, following the above-described methodology. We quantified the protein concentrations using the Pierce™ 660 nm Protein Assay Reagent, with bovine serum albumin serving as the standard.

### Toggling with dual binary inputs

DH10β Δ*cyaA* cells harboring the F1RP*-GFP/F2P*/SP were first grown under permissive conditions (0.5 mM BzF at 30 °C, binary code = 11) for 24 h. Post this incubation, the cell density (OD_600_) and sfGFP fluorescence were measured to assess the level of the gene of interest (GOI) expression. The cultures were then centrifuged at 4 °C and 3,000 g for 10 min and subjected to three washing steps with LB. Following washing, the cultures were diluted 100-fold in fresh LB broth, supplemented with Amp, Kan, Spec, and IPTG, and shifted to either of three non-permissive conditions, characterized by the binary codes 00, 01, or 10, and incubated for an additional 24 h. After each switch, cultures were incubated for 24 h, and then the level of GOI expression was assessed through sfGFP fluorescence measurement. This alternating pattern of shifting between permissive and non-permissive conditions was repeated for a total of two cycles to comprehensively gauge the system’s behavior.

### Input switching

DH10β Δ*cyaA* cells harboring the F1RP*-GFP/F2P*/SP were cultivated for 24 h under non-permissive conditions (no BzF at 37 °C, binary code = 00). Subsequently, they were diluted 200-fold into two distinct non-permissive media (binary codes 01 and 10). To induce sfGFP expression, a transition to the permissive conditions (11) was initiated. This was accomplished either by adding BzF (0.5 mM) to the media or by decreasing the incubation temperature from 37 °C to 30 °C at set intervals: 3, 6, or 9 hpi. After a total incubation period of 24 h, the fluorescence intensities of the bacterial cultures were assessed using the SpectraMax iD3 Multi-Mode Microplate Reader and subsequently normalized to the OD_600_ measurements.

### Determination of doubling time

Strains were pre-cultivated at 37 °C for 24 h. After this initial growth phase, the culture was diluted 200-fold into LB broth and then incubated under the designated culture conditions with constant shaking at 300 rpm for 24 h. Every 30 minutes, the OD_600_ was recorded using a SpectroStar Nano Microplate Reader (BMG Labtech, Germany). The doubling time for each strain was deduced from the exponential growth phase data using the exponential growth equation in GraphPad Prism 10. Only datasets with an R-squared value greater than 0.95 were considered for accurate doubling time estimation.

## Data availability

Supplementary figures and tables are available in the online supplementary information.

## Acknowledgements

We thank Prof. Peter G. Schultz for providing the pKTECM, pKTECB, pT25SC-rep, pBK-SCT18, and pUltra-BzF plasmids and Prof. Anders Løbner-Olesen for providing the T18/T25 fusion constructs. This work was supported by the National Research Foundation of Korea (NRF) grant funded by the Korea government (MSIT) [2021R1C1C1005766 and 2021R1A4A1021950].

## Author contributions

M.K. conceived and supervised the project. J.C., J.A., J.B., and M.Y. performed the experiments and analyzed the data. J.C., H.Y., and M.K. wrote the paper with input from all authors.

