## Supplementary Figures and Tables for "Impact of exogenous aminoacyl-tRNA synthetase and tRNA on temperature sensitivity in *Escherichia coli*"

### 1. Supplementary Figures

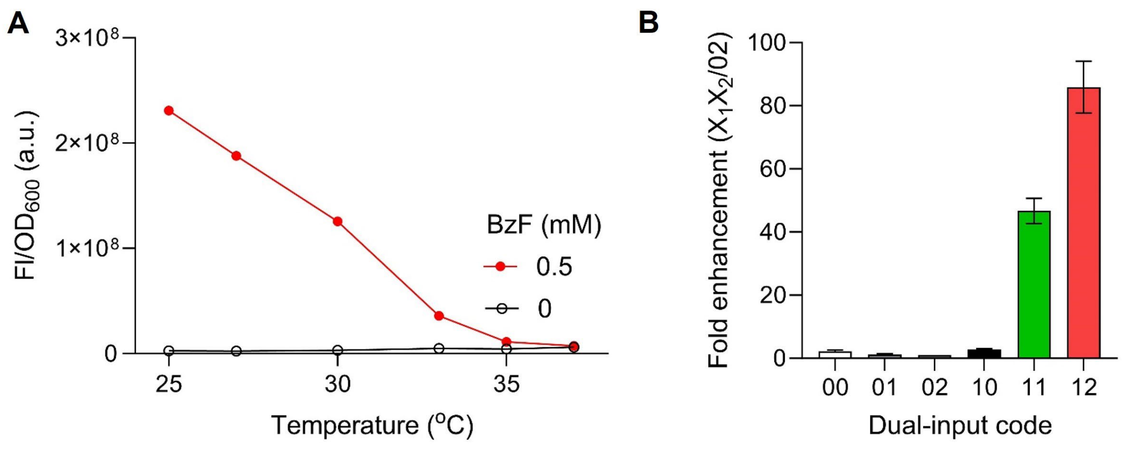

**Supplementary Fig. 1 Characterization of a dual-input biologic gate. A** Normalized super-folder green fluorescent protein (sfGFP) fluorescence signals were plotted across temperatures (25 °C, 27 °C, 30 °C, 33 °C, 35 °C, and 37 °C) after incubating DH10β Δ*cyaA*/F1RP*-GFP/F2P*/SP cells in the specified conditions in the absence or presence of BzF (0.5 mM) for 24 h. **B** Relative sfGFP expression was compared across six conditions (Code $X_1X_2$ = 00, 01, 10, 11, 02, and 12). sfGFP fluorescence intensities were normalized to the non-permissive condition (Code $X_1X_2$ = 02). Error bars represent standard deviation (n = 4). The codes $X_1X_2$ refer to BzF concentrations (0 mM, $X_1$ = 0; 0.5 mM, $X_1$ = 1) and temperature conditions (37 °C, $X_2$ = 0; 30 °C, $X_2$ = 1; 25 °C, $X_2$ = 2). Fl, fluorescence; and $OD_{600}$, optical density at 600 nm.

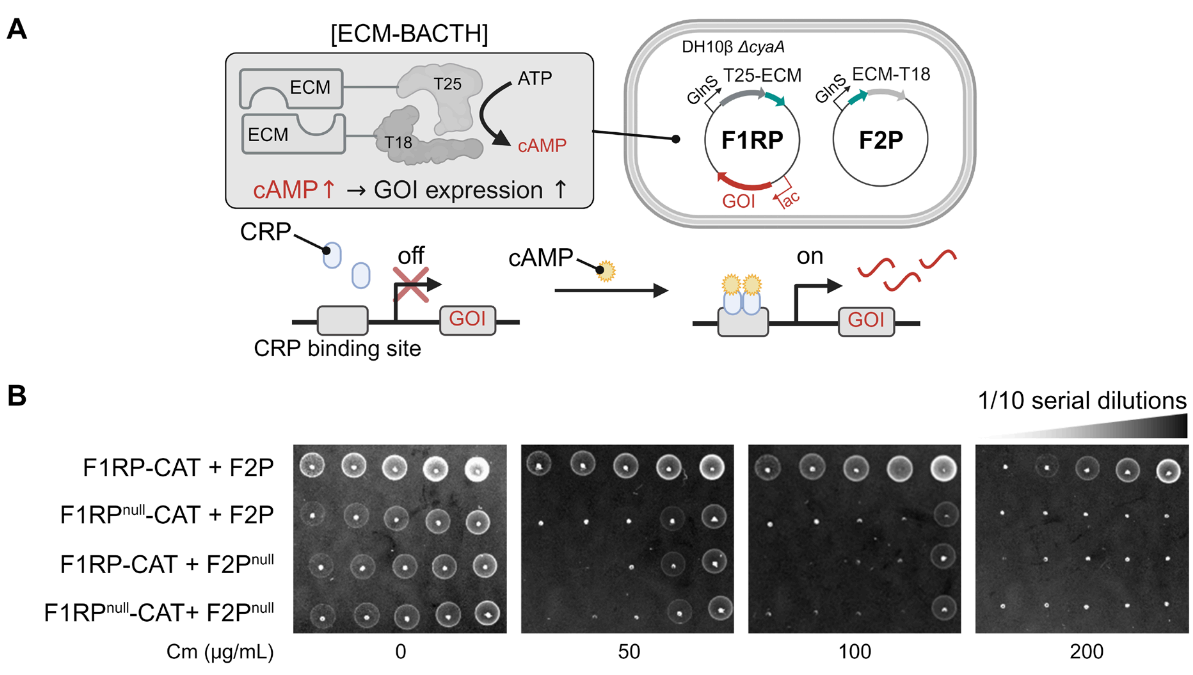

**Supplementary Fig. 2 An *E. coli* chorismate mutase (ECM) mediated bacterial adenylate cyclase two-hybrid system (ECM-BACTH). A** Schematic representation of the ECM-BACTH system. In this system, fusion plasmids F1RP and F2P are engineered to express T25-ECM and ECM-T18 fusion proteins, respectively. ECM dimerization triggers T18-T25 dimerization, leading to the production of cAMP. As the DH10β Δ*cyaA* strain lacks cAMP, the expression of the GOI, controlled by the cAMP-dependent *lac* promoter, is reliant on ECM dimerization. **B** Dilution spot images of the DH10β Δ*cyaA* cells harboring F1RP-CAT/F2P, F1RP$^{null}$-CAT/F2P, F1RP-CAT/F2P$^{null}$, or F1RP$^{null}$-CAT/ F2P$^{null}$ on LB-agar plates, both in the absence and presence of chloramphenicol (Cm, 50, 100, and 200 μg/mL), incubated at 37 °C for 20 h. 'F1RP-CAT' represents the T25-ECM fusion plasmid containing the cat gene, and 'F2P' refers to the ECM-T18 fusion plasmid. For control experiments, ECM genes were deleted from F1RP and F2P to yield 'F1RP$^{null}$-CAT' and 'F2P$^{null}$', respectively. CRP refers to cAMP receptor protein and CAT represents chloramphenicol acetyltransferase.

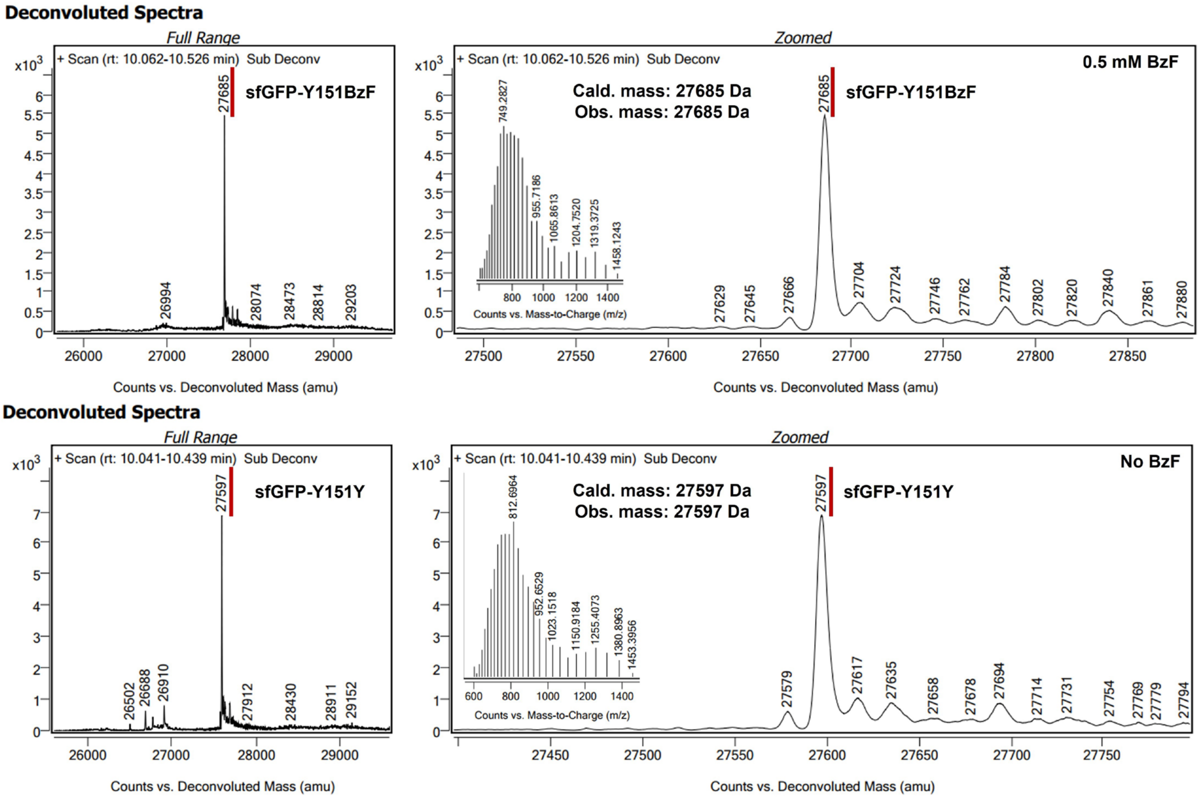

**Supplementary Fig. 3 Mass spectra of sfGFP variants obtained from the culture of DH10β/pET22b-T5-sfGFP*/SP in the presence or absence of BzF.**

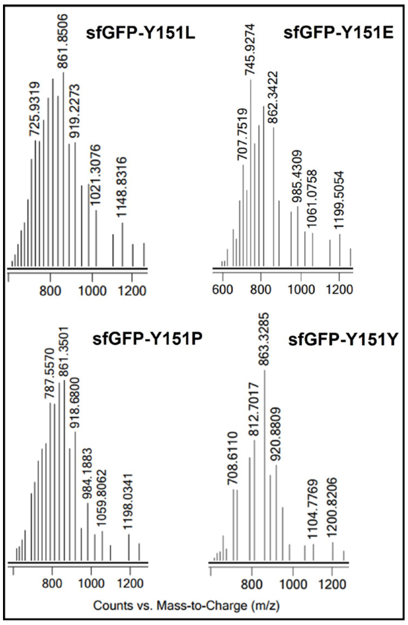

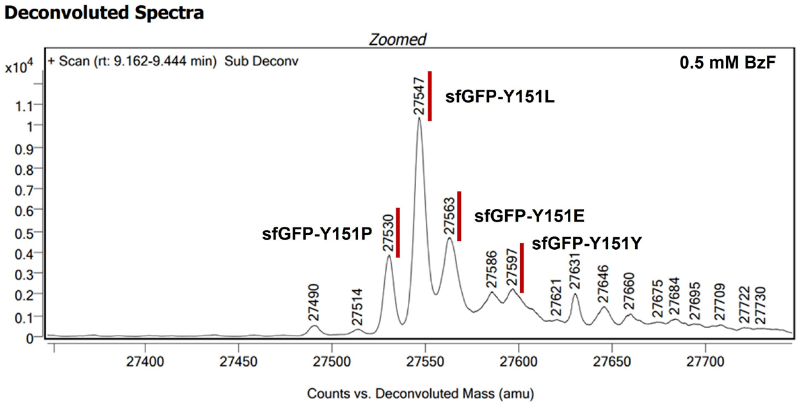

| Amino acid residue at 151 site | Leu | Glu | Pro | Tyr |
|---|---|---|---|---|
| Cald. mass (Da) | 27547 | 27563 | 27531 | 27597 |
| Obs. mass (Da) | 27547 | 27563 | 27530 | 27597 |

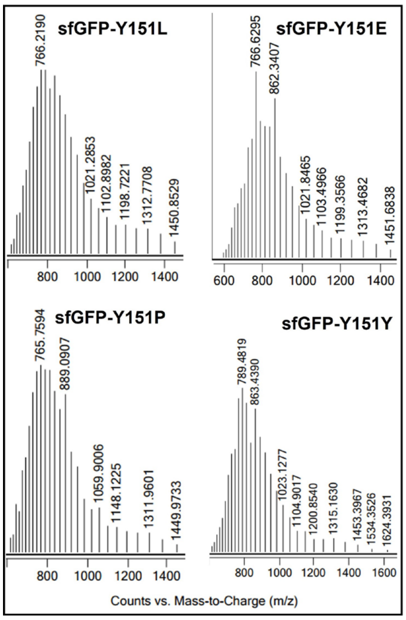

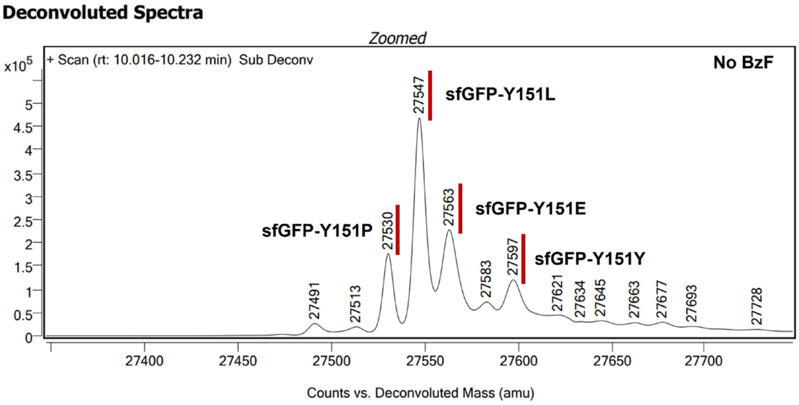

| Amino acid residue at 151 site | Leu | Glu | Pro | Tyr |
|---|---|---|---|---|
| Cald. mass (Da) | 27547 | 27563 | 27531 | 27597 |
| Obs. mass (Da) | 27547 | 27563 | 27530 | 27597 |

**Supplementary Fig. 4 Mass spectra of sfGFP variants obtained from the culture of DH10β/pET22b-T5-sfGFP*/SP-aaRS$^{null}$ in the presence or absence of BzF.**

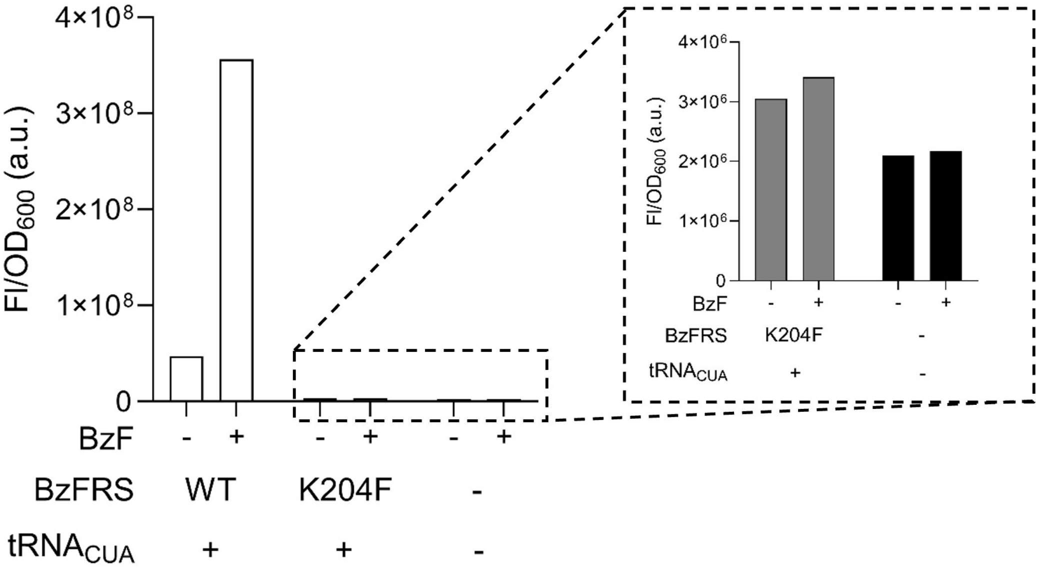

**Supplementary Fig. 5 Comparison of amber suppression efficiency of BzFRS/tRNA$_{CUA}$ and BzFRS-K204F/tRNA$_{CUA}$.** The expression of sfGFP* gene, which contains an amber (TAG) codon at position 151, was assessed under defined conditions through sfGFP fluorescence. The results for BzFRS-K204F and the null variant were magnified for clarity.

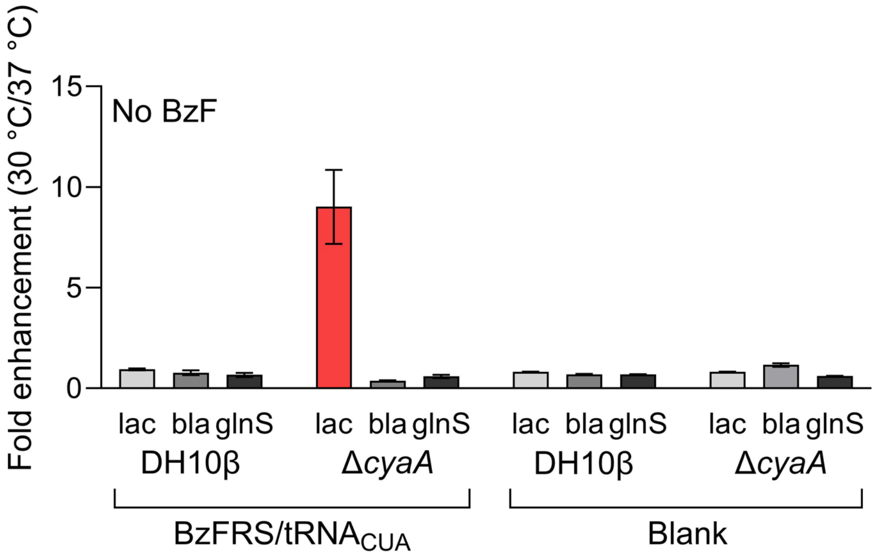

**Supplementary Fig. 6 *Lac* promoter-specific temperature sensitivity.** The enhancement in gene expression levels at 30 °C compared to 37 °C, controlled by three different promoters (*lac*, *bla*, and *glnS*), was assessed both with and without BzFRS/$tRNA_{CUA}$ in the DH10β and DH10β Δ*cyaA* strains grown in LB. The data presented is a ratio of fluorescence intensities at 30 °C and 37 °C. n = 4 and error bars stand for standard deviation.

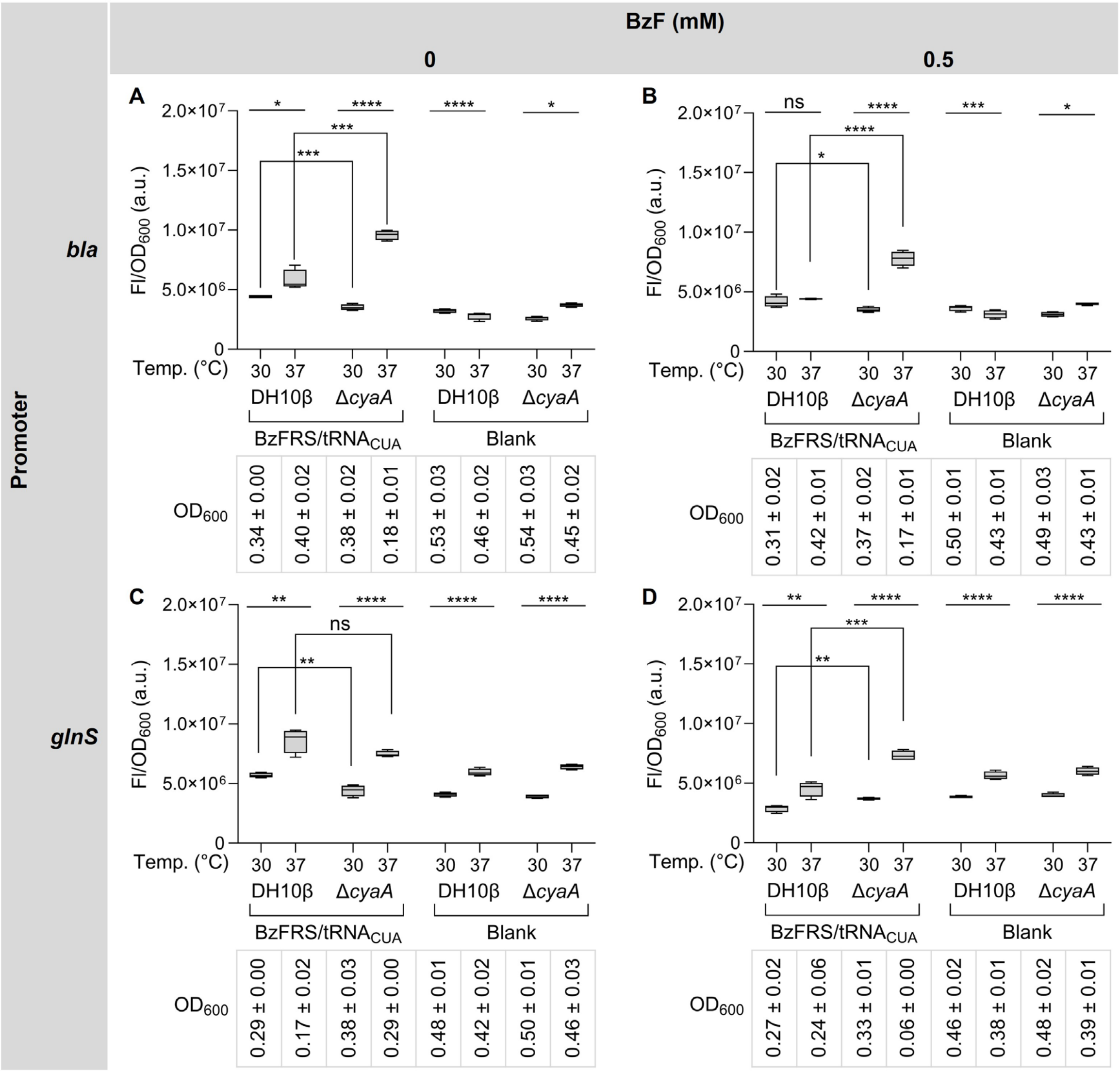

**Supplementary Fig. 7 Impact of exogenous BzFRS and $tRNA_{CUA}$ on temperature sensitivity in gene expression under the control of the *bla* or *glnS* promoter.** DH10β and DH10β Δ*cyaA* cells harboring F1RP-GFP-bla/F2P/SP, F1RP-GFP-bla/F2P/$SP^{null}$ (**A**, **B**), F1RP-GFP-glnS/F2P/SP, or F1RP-GFP-glnS/F2P/$SP^{null}$ (**C**, **D**) plasmids were incubated in LB broth in the absence (**A**, **C**) and presence (**B**, **D**) of BzF (0.5 mM) under specified temperatures for 24 h and the expression levels of the sfGFP were quantified by fluorescence reading (Ex/Em: 485 nm/525 nm). n = 4; box limits indicate the interquartile range; whiskers represent the range from minimum to maximum; the center line denotes the median; ns (not significant), * P < 0.05, ** P < 0.01, *** P < 0.001, **** P < 0.0001 by Student's t-test; and variability in $OD_{600}$ values is represented by the standard deviation.

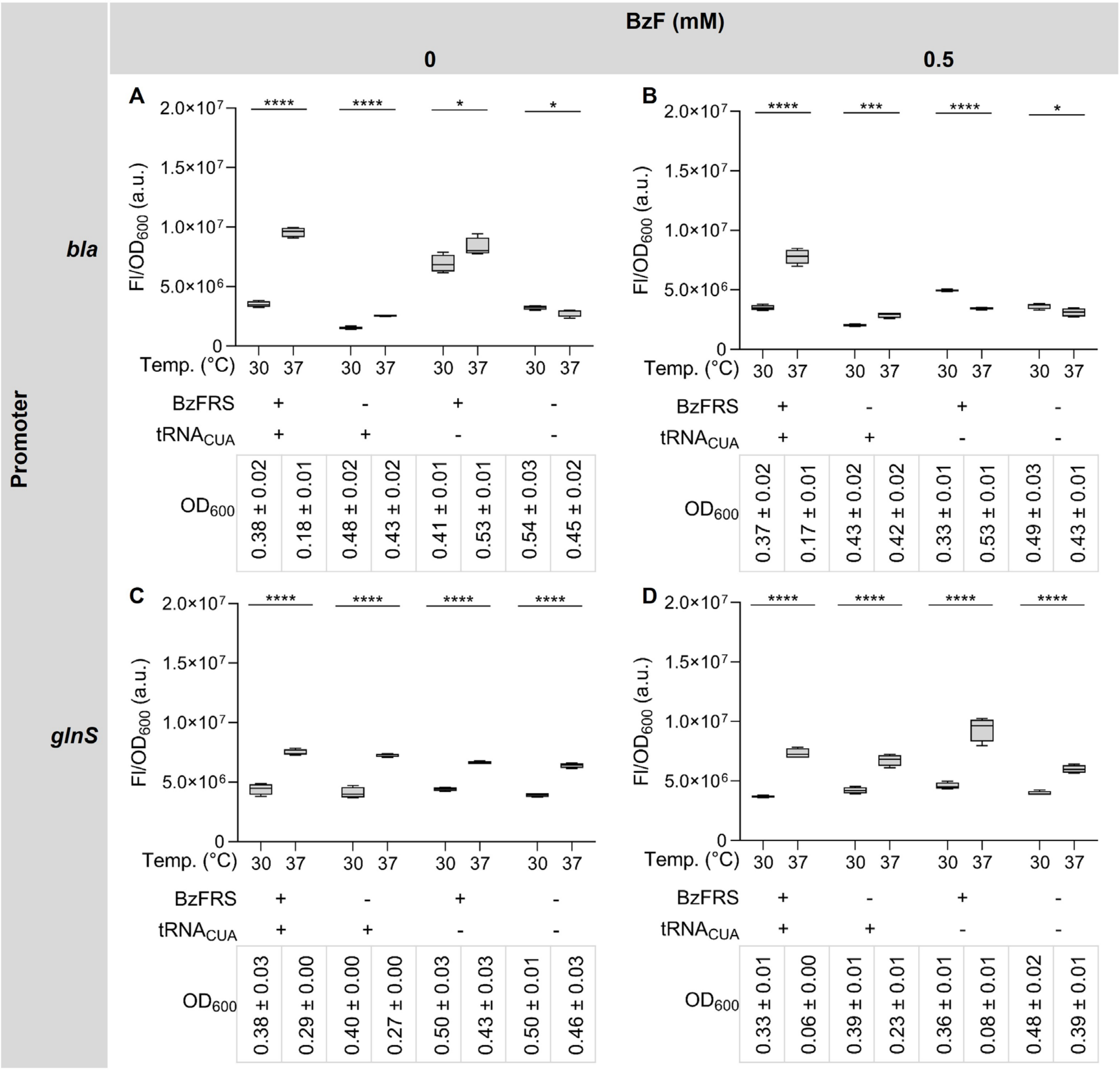

**Supplementary Fig. 8 Assessing the impact of exogenous BzFRS or tRNA$_{CUA}$ on gene expression controlled by the *bla* or *glnS* promoter.** The influence of either SP, SP-aaRS$^{null}$, SP-tRNA$^{null}$, or SP$^{null}$ on sfGFP expression was assessed in DH10β Δ*cyaA*/F1RP-GFP-bla/F2P (**A**, **B**) and DH10β Δ*cyaA*/F1RP-GFP-glnS/F2P (**C**, **D**). This assessment was carried out in the absence (**A**, **C**) and presence (**B**, **D**) of BzF (0.5 mM) at specified temperatures. Cells were incubated in LB for 24 h before measuring fluorescence. n = 4; box limits indicate the interquartile range; whiskers represent the range from minimum to maximum; the center line denotes the median; and * $P < 0.05$, *** $P < 0.001$, **** $P < 0.0001$ by Student's t-test; and variability in $OD_{600}$ values is represented by the standard deviation.

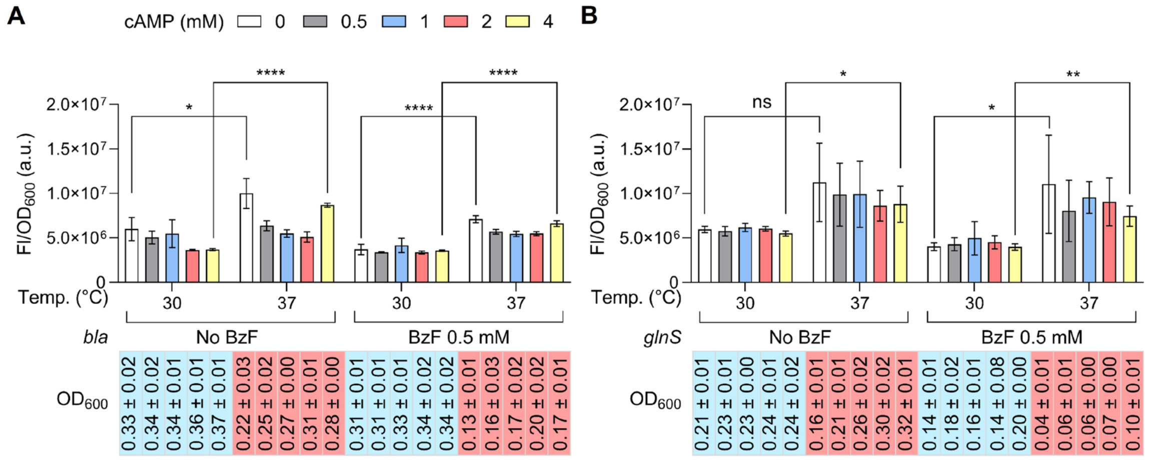

**Supplementary Fig. 9 Evaluating the impact of cAMP addition on gene expression controlled by the *bla* or *glnS* promoter.** The sfGFP fluorescence was assessed after incubating DH10β Δ*cyaA*/F2P/SP cells containing either the F1RP-GFP-bla (**A**) or F1RP-GFP-glnS (**B**) plasmids under various cAMP concentrations (0, 0.5, 1, 2, and 4 mM) at both 30 °C and 37 °C. Cells were incubated in LB broth both in the absence and presence of BzF (0.5 mM) for 24 h. n = 4; ns (not significant), * $P < 0.05$, ** $P < 0.01$, **** $P < 0.0001$ by Student's t-test; error bars represent the standard deviation; and variability in $OD_{600}$ values is represented by the standard deviation.

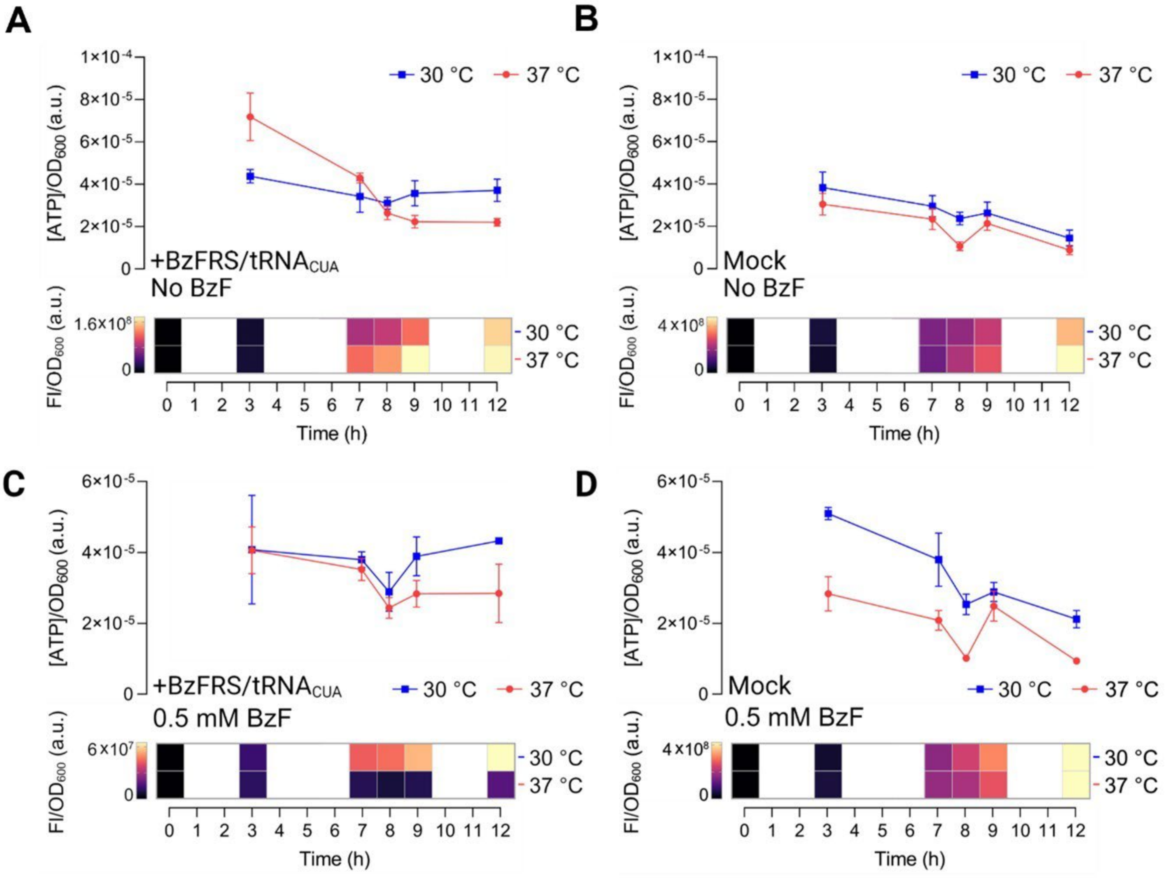

**Supplementary Fig. 10 Time-course measurement of intracellular ATP and sfGFP fluorescence.** The intracellular ATP levels and sfGFP fluorescence of DH10β Δ*cyaA*/F1RP-GFP/F2P cells harboring either the SP (**A**, **C**) or SP$^{null}$ (**B**, **D**) plasmid were assessed. Cells were incubated in LB broth in the absence (**A**, **B**) and presence (**C**, **D**) of BzF (0.5 mM) for 12 h at either 30 °C or 37 °C. Measurements for ATP levels and sfGFP fluorescence were taken at specific intervals: 3 h, 7 h, 8 h, 9 h, and 12 h. For each condition, n = 4; error bars represent the standard deviation.

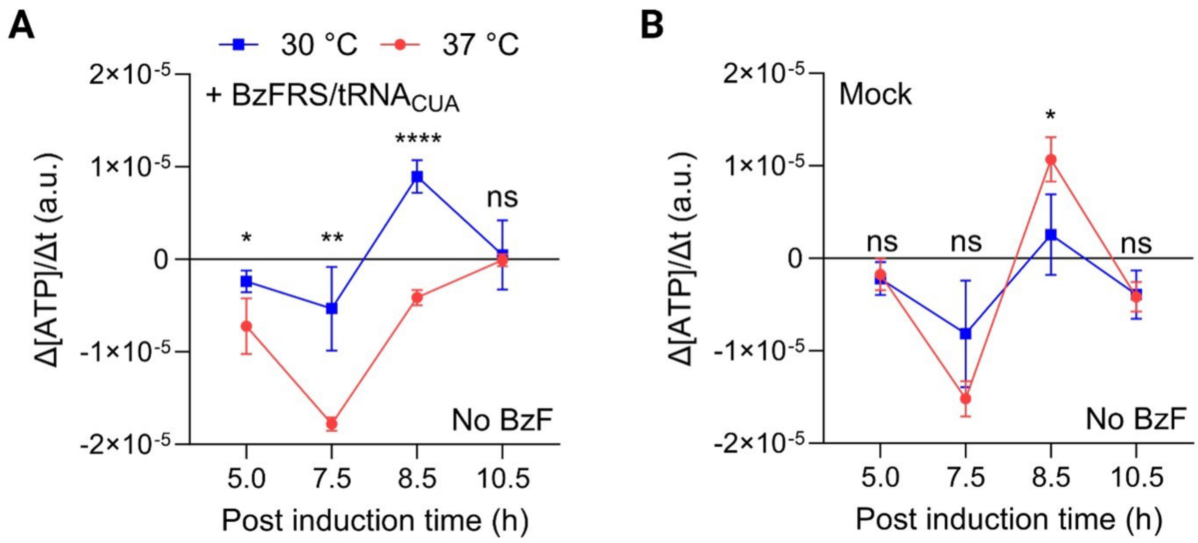

**Supplementary Fig. 11 Changes in ATP concentration slopes over time.** The rate of intracellular ATP changes in DH10β Δ*cyaA*/F1RP-GFP/F2P cells harboring either the SP (**A**) or $SP^{null}$ (**B**) plasmid was assessed. Cells were incubated in LB broth in the absence of BzF for 12 h at either 30 °C or 37 °C. ATP slopes over time (Δ[ATP]/Δt) were calculated and plotted at specific intervals: 5 h, 7.5 h, 8.5 h, and 10.5 h. For each condition, n = 4; ns (not significant), * $P < 0.05$, ** $P < 0.01$, **** $P < 0.0001$ by Student's t-test; error bars represent the standard deviation.

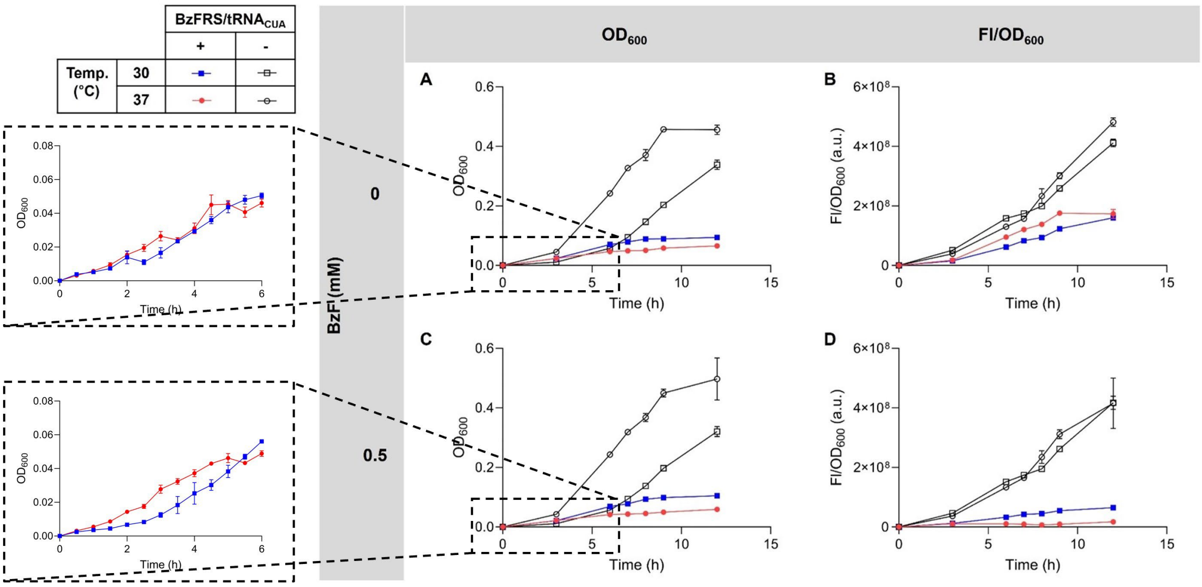

**Supplementary Fig. 12 Growth curve and time-course measurement of sfGFP expression.** The $OD_{600}$ and sfGFP fluorescence of DH10β Δ*cyaA*/F1RP-GFP/F2P cells harboring either the SP or $SP^{null}$ plasmid were recorded. Cells were incubated in LB broth both in the absence and presence of BzF (0.5 mM) for 12 h at either 30 °C or 37 °C. Measurements for $OD_{600}$ (**A**, **C**) and sfGFP fluorescence (**B**, **D**) were taken at specific intervals: 3 h, 6 h, 7 h, 8 h, 9 h, and 12 h. For each condition, n = 4; error bars represent the standard deviation. The early growth phase is highlighted in the left panel, where additional data points are provided for greater detail.

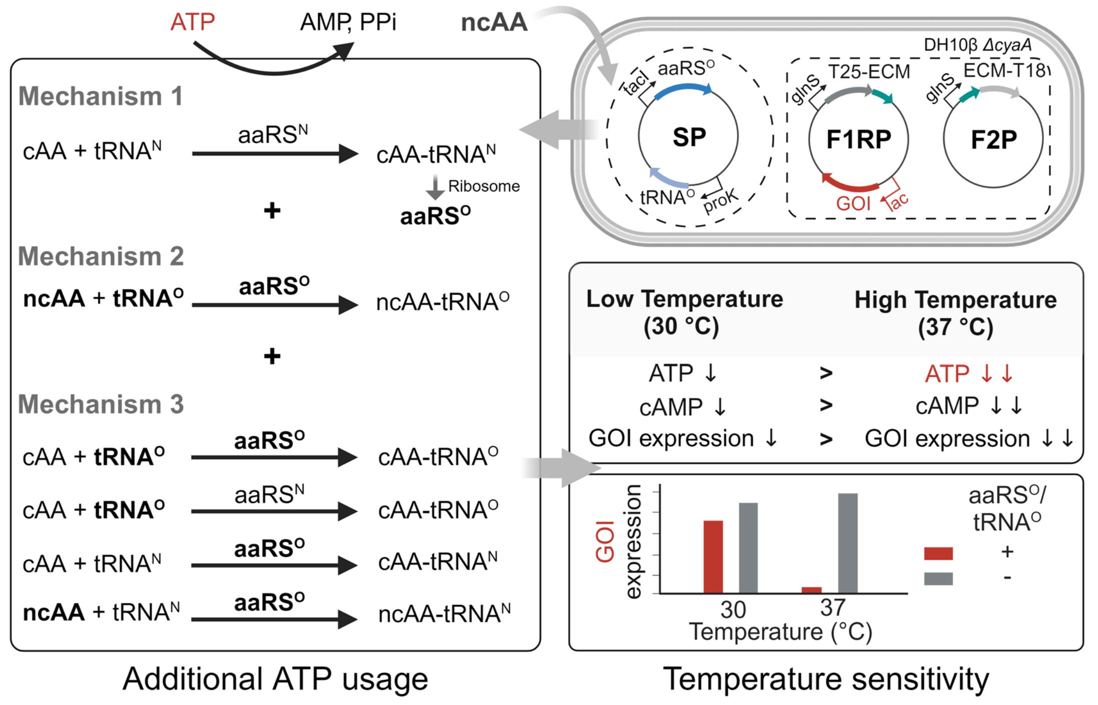

**Supplementary Fig. 13 Mechanistic overview of temperature sensitivity in a bacterial adenylate cyclase two-hybrid (BACTH) system caused by exogenous orthogonal aminoacyl-tRNA synthetase (aaRS$^{O}$) and orthogonal tRNA (tRNA$^{O}$).** The fusion plasmids F1RP and F2P complement cAMP deficiency in the *E. coli ΔcyaA* strain by expressing T25-ECM and ECM-T18, respectively. The increased consumption of intracellular ATP can be caused by inherent aminoacylation process (Mechanism 1) for the overexpression of exogenous genes. Additionally, overexpression of aaRS$^{O}$ and tRNA$^{O}$ encoded in the suppressor plasmid SP leads to additional consumption of intracellular ATP at high temperatures (Mechanism 2). Furthermore, in instances where aaRS$^{O}$ and/or tRNA$^{O}$ are not perfectly orthogonal, unintended cross-reactions with native aaRS (aaRS$^{N}$) or tRNA (tRNA$^{N}$) can also cause ATP depletion (Mechanism 3). Such reactions can utilize canonical amino acids (cAAs) and proceeds even in the absence of non-canonical amino acids (ncAAs). The resultant depletion of starting materials triggers a shortage of cAMP and hampers the expression of the gene of interest (GOI) at high temperatures. ECM, wild type *E. coli* chorismate mutase; and SP refers to suppressor plasmid.

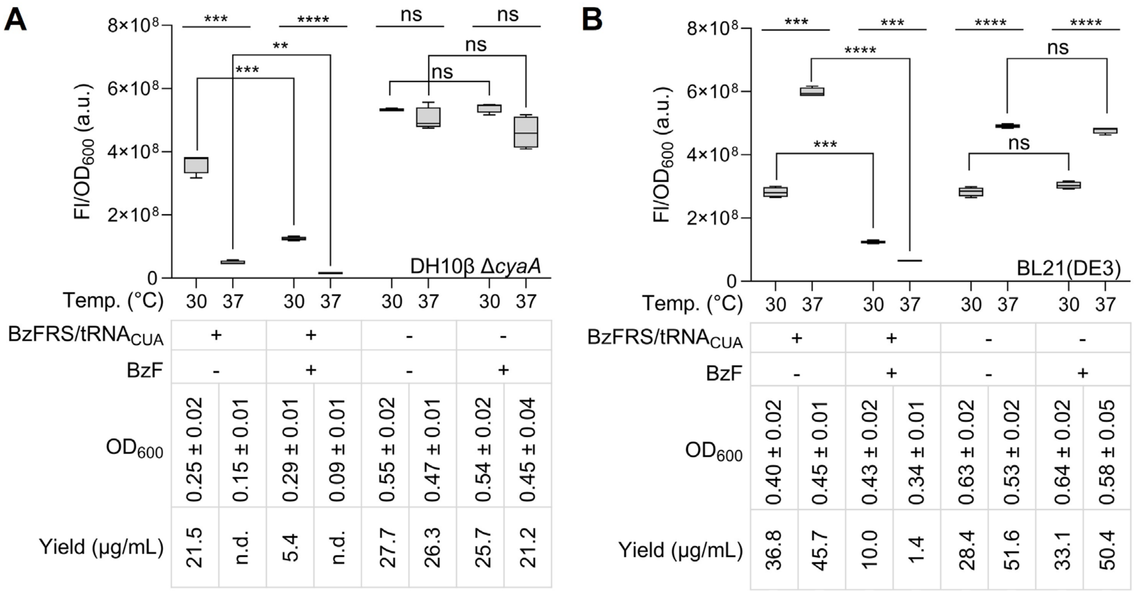

**Supplementary Fig. 14 Comparison of sfGFP expression in DH10β Δ*cyaA* and BL21(DE3).** The $OD_{600}$ and sfGFP fluorescence were measured for each bacterial strain under specified conditions. Following purification, the yield was calculated using bovine serum albumin as a standard. FI/$OD_{600}$ data are shown in the upper panel, and the $OD_{600}$ and the purified sfGFP yields are summarized in the lower panel for the cultures of DH10β Δ*cyaA* (**A**) and BL21(DE3) (**B**). n = 4; box limits indicate the interquartile range; whiskers represent the range from minimum to maximum; the center line denotes the median; ns (not significant), ** $P < 0.01$, *** $P < 0.001$, **** $P < 0.0001$ by Student's t-test; variability in $OD_{600}$ values is represented by the standard deviation; the purification yields are derived from a representative set; n.d. means not determined.

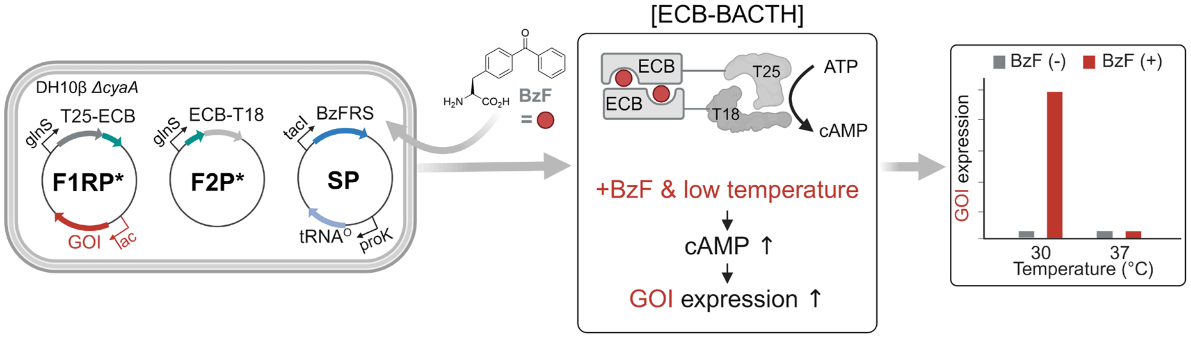

**Supplementary Fig. 15 The ECM was replaced with its BzF dependent variant ECB to afford ECB-BACTH system.** F1RP*, T25-ECB fusion plasmid containing reporter gene; F2P*, ECB-T18 fusion plasmid; and SP, suppressor plasmid.

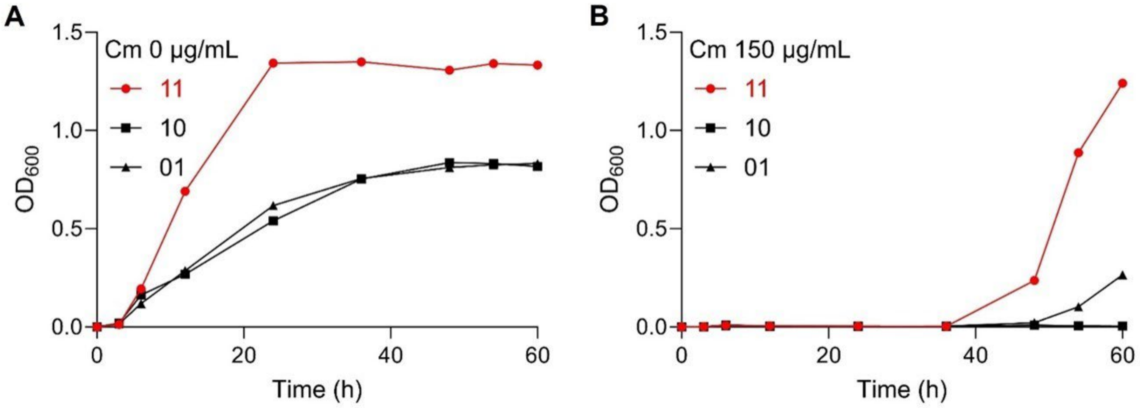

**Supplementary Fig. 16 Growth curves of the DH10β Δ*cyaA* with ECB-BACTH.** Growth curves of DH10β Δ*cyaA*/F1RP*-CAT/F2P*/SP cells were examined under various culture conditions, both in the absence (**A**) and presence (**B**) of Cm (150 μg/mL). The binary codes 11, 10, and 01 correspond to conditions [0.5 mM BzF and 30 °C], [0.5 mM BzF and 37 °C], and [No BzF and 30 °C], respectively.

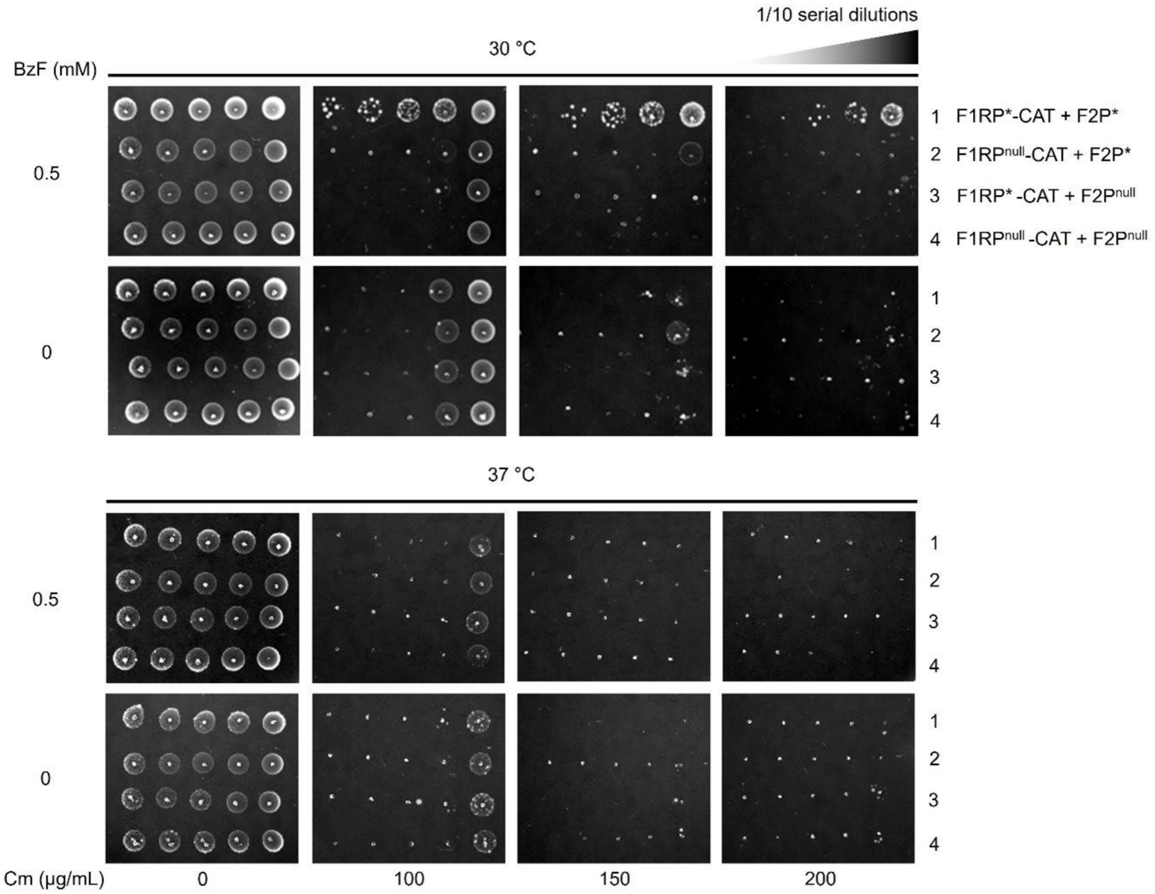

**Supplementary Fig. 17 Growth of DH10β Δ*cyaA* with ECB-BACTH.** Dilution spot images show DH10β Δ*cyaA*/SP cells harboring F1RP*-CAT/F2P*, F1RP$^{null}$-CAT/F2P*, F1RP*-CAT/F2P$^{null}$, or F1RP$^{null}$-CAT/F2P$^{null}$ on LB-agar plates. These were grown with or without Cm (100, 150, and 200 μg/mL) under specified conditions: presence/absence of 0.5 mM BzF and at either 30 °C or 37 °C for 72 h.

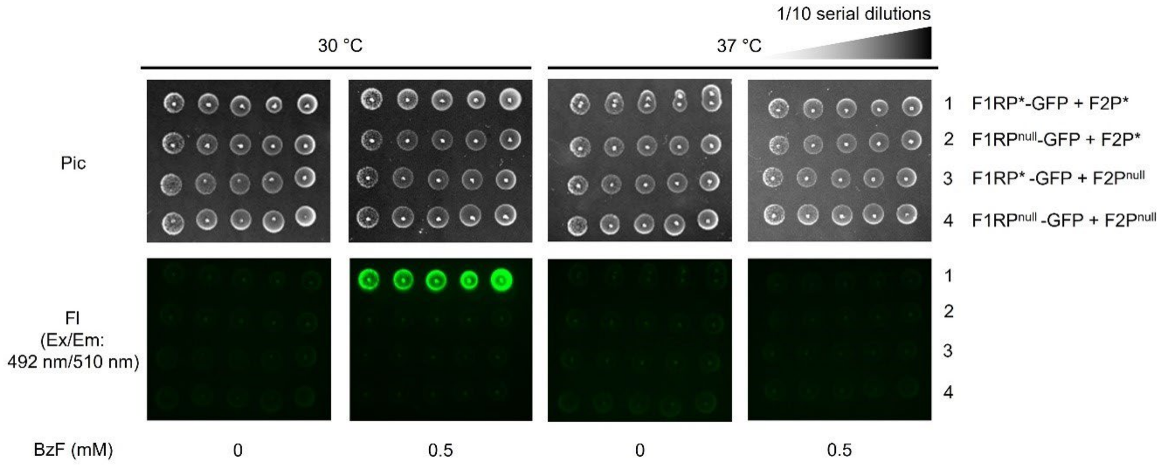

**Supplementary Fig. 18 Fluorescence imaging of DH10β Δ*cyaA* with ECB-BACTH.** Dilution spot images (Pic) of the DH10β Δ*cyaA*/SP cells harboring F1RP*-GFP/F2P*, F1RP$^{null}$-GFP/F2P*, F1RP*-GFP/F2P$^{null}$, or F1RP$^{null}$-GFP/F2P$^{null}$ on LB-agar. Concurrently, sfGFP fluorescence (FI) was assessed. Cells were incubated under specific conditions: with or without 0.5 mM BzF and at either 30 °C or 37 °C for 72 h.

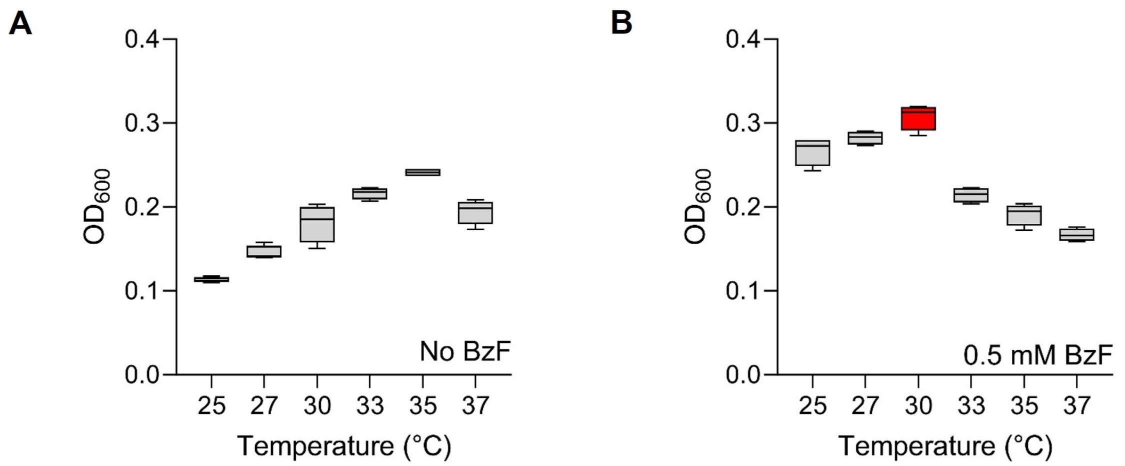

**Supplementary Fig. 19 Growth profile of DH10β Δ*cyaA* with ECB-BACTH at different temperatures.** Optical densities at 600 nm ($OD_{600}$) were plotted across temperatures (25 °C, 27 °C, 30 °C, 33 °C, 35 °C, and 37 °C) after incubating DH10β Δ*cyaA*/F1RP*-GFP/F2P*/SP cells in the specified conditions in the absence (**A**) or presence (**B**) of BzF (0.5 mM) for 24 h. n = 4; box limits indicate the interquartile range; whiskers represent the range from minimum to maximum; and the center line denotes the median.

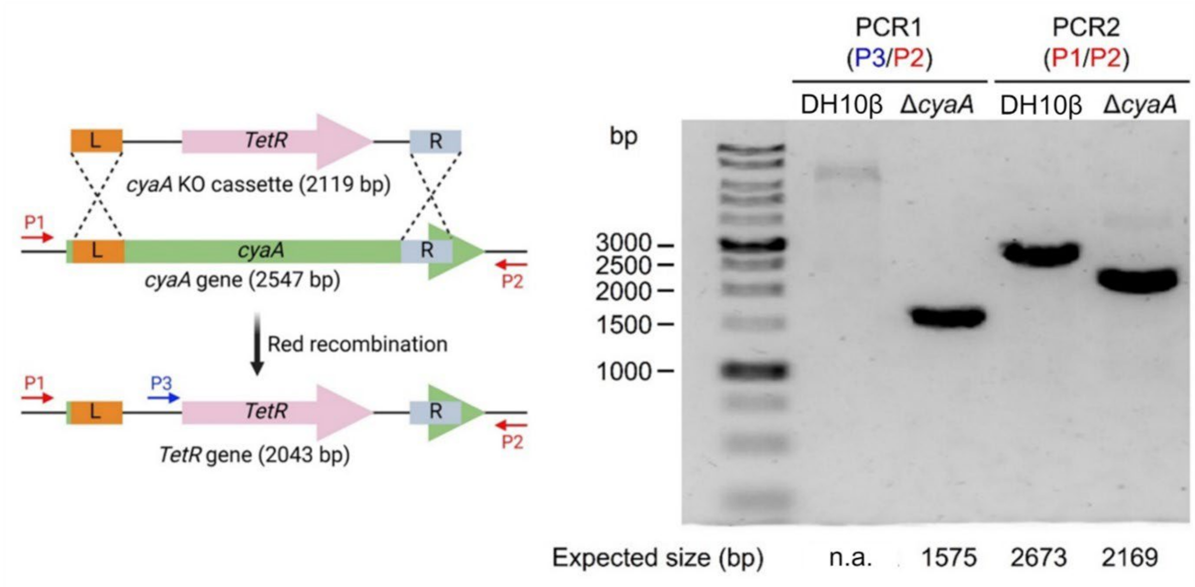

**Supplementary Fig. 20 Verification of *cyaA* knockout in DH10β.** Colony PCR products from either DH10β or DH10β Δ*cyaA*, using P3/P2 or P1/P2 primers, were analyzed by electrophoresis on a 1% agarose gel. P1: oMK-58, P2: oMK-59, and P3: oMK-28.

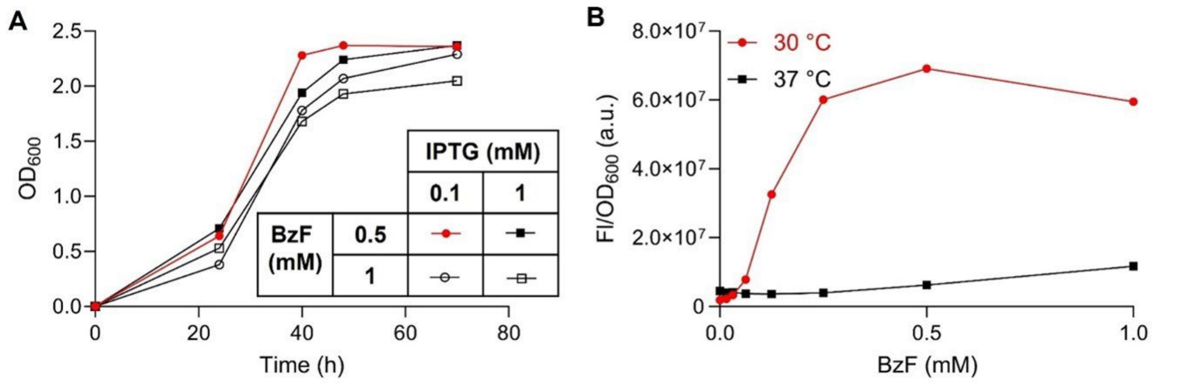

**Supplementary Fig. 21 Screening for optimal concentration of IPTG and BzF. A** Growth curve of DH10β Δ*cyaA*/F1RP*-CAT/F2P*/SP cells at 30 °C under various culture conditions: LB supplemented with Cm (50 μg/mL) and combinations of IPTG (either 0.1 mM or 1 mM) and BzF (either 0.5 mM or 1 mM). **B** The normalized sfGFP fluorescence was plotted against BzF concentrations (0, 0.0156, 0.0313, 0.0625, 0.125, 0.25, 0.5, and 1 mM) after incubating DH10β Δ*cyaA*/F1RP*-GFP/F2P*/SP cells for 24 h at two temperature conditions (30 °C and 37 °C).

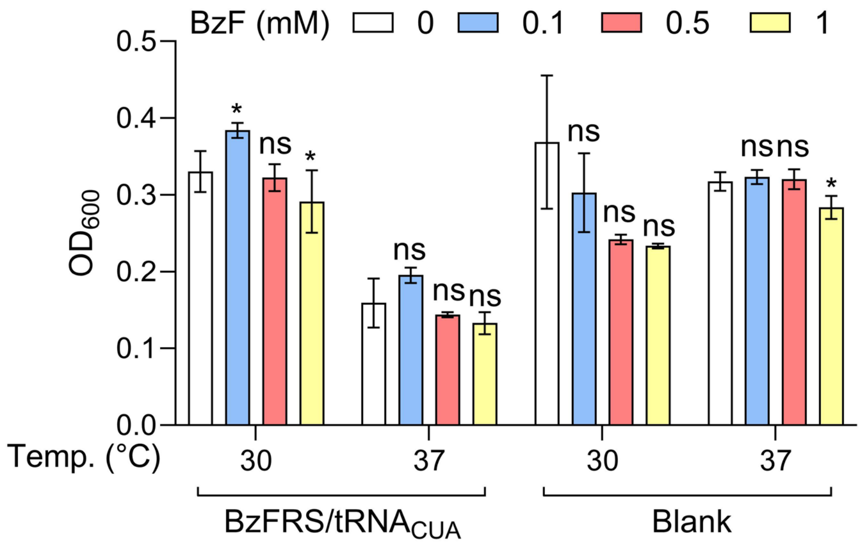

**Supplementary Fig. 22 Evaluation of the impact of BzF addition on growth.** The $OD_{600}$ was assessed for DH10β Δ*cyaA*/F1RP-CAT/F2P cells harboring either the SP or $SP^{null}$ plasmid, incubated in LB broth at various BzF concentrations (0, 0.1, 0.5, and 1 mM) and temperatures (30 °C and 37 °C) over 24 h. n = 4; ns (not significant), * $P < 0.05$ by Student's t-test; and error bars represent the standard deviation.

### 2. Supplementary Tables

**Supplementary Table 1.** Plasmid used in this study.

| Plasmid Name | Marker | Origin | ORF1 | | ORF2 | | Reference |
|---|---|---|---|---|---|---|---|
| | | | Promoter | Gene | Promoter | Gene | |
| pKTECM | $kan^R$ | ColE1 | $P_{tet}$ | ECM | | | [1] |
| pKTECB | $kan^R$ | ColE1 | $P_{tet}$ | ECB | | | [1] |
| pT25SC-rep | $amp^R$ | P15A | $P_{glnS}$ | T25-SC | $P_{lac}$ | CAT | [2] |
| pBK-SCT18 | $kan^R$ | ColE1 | $P_{glnS}$ | SC-T18 | | | [2] |
| pKD4-$tet^R$ | $amp^R$ | R6Kγ | | $tet^R$ | | | [3] |
| pKD46 | $amp^R$ | pSC101 | $P_{BAD}$ | γ, β, exo | | rep101 ts | [3] |
| pET22b-T5-sfGFP | $amp^R$ | ColE1 | $P_{T7}$-$P_{T5}$ | sfGFP | | | [4] |
| pET22b-T5-sfGFP* | $amp^R$ | ColE1 | $P_{T7}$-$P_{T5}$ | sfGFP-Y151am | | | [5] |
| pET22b-T5-Myo | $amp^R$ | ColE1 | $P_{T7}$-$P_{T5}$ | Sperm whale myoglobin | | | [6] |
| F1RP-CAT | $amp^R$ | P15A | $P_{glnS}$ | T25-ECM | $P_{lac}$ | CAT | This study |
| F1RP-GFP | $amp^R$ | P15A | $P_{glnS}$ | T25-ECM | $P_{lac}$ | sfGFP | This study |
| F1RP-GFP-bla | $amp^R$ | P15A | $P_{glnS}$ | T25-ECM | $P_{bla}$ | sfGFP | This study |
| F1RP-GFP-glnS | $amp^R$ | P15A | $P_{glnS}$ | T25-ECM | $P_{glnS}$ | sfGFP | This study |
| F1RP*-CAT | $amp^R$ | P15A | $P_{glnS}$ | T25-ECB | $P_{lac}$ | CAT | This study |
| F1RP*-GFP | $amp^R$ | P15A | $P_{glnS}$ | T25-ECB | $P_{lac}$ | sfGFP | This study |
| $F1RP^{null}$-CAT | $amp^R$ | P15A | $P_{glnS}$ | T25 | $P_{lac}$ | CAT | This study |
| $F1RP^{null}$-GFP | $amp^R$ | P15A | $P_{glnS}$ | T25 | $P_{lac}$ | sfGFP | This study |
| F2P | $kan^R$ | ColE1 | $P_{glnS}$ | ECM-T18 | | | This study |
| F2P* | $kan^R$ | ColE1 | $P_{glnS}$ | ECB-T18 | | | This study |
| $F2P^{null}$ (pBK-T18-Blk) | $kan^R$ | ColE1 | $P_{glnS}$ | T18 | | | [2] |
| SP (pUltra-BzF) | $spec^R$ | CDF | $P_{tacI}$ | *Mj*BzFRS | $P_{proK}$ | *Mj*$tRNA_{CUA}$ | [1] |
| SP-$tRNA^{null}$ | $spec^R$ | CDF | $P_{tacI}$ | *Mj*BzFRS | | | This study |
| SP-$aaRS^{null}$ | $spec^R$ | CDF | $P_{proK}$ | *Mj*$tRNA_{CUA}$ | | | This study |
| $SP^{null}$ | $spec^R$ | CDF | | | | | This study |
| $SP^{null}$-Myo | $spec^R$ | CDF | $P_{tacI}$ | Sperm whale myoglobin | | | This study |
| SP-K204F | $spec^R$ | CDF | $P_{tacI}$ | *Mj*BzFRS-K204F | $P_{proK}$ | *Mj*$tRNA_{CUA}$ | This study |
| SP-K204F-$tRNA^{null}$ | $spec^R$ | CDF | $P_{tacI}$ | *Mj*BzFRS-K204F | | | This study |

**Supplementary Table 2.** Mass spectrometry data of sfGFP variants.

| Exogenous BzFRS/$tRNA_{CUA}$ | BzF (mM) | Cald. Mass, (Da) | Obs. Mass, (Da) | Amino Acid Residue at 151am Site | Yield (mg/L) |
|---|---|---|---|---|---|
| BzFRS/$tRNA_{CUA}$ | 0 | 27,597 | 27,597 | Tyr | 0.17 |
| | 0.5 | 27,685 | 27,685 | BzF | 8.97 |
| $tRNA_{CUA}$ | 0 or 0.5 | 27,547 | 27,547 | Leu | 0.18 (BzF 0 mM)<br>0.11 (BzF 0.5 mM) |
| | | 27,563 | 27,563 | Glu | |
| | | 27,531 | 27,530 | Pro | |
| | | 27,597 | 27,597 | Tyr | |
| No BzFRS/$tRNA_{CUA}$ | 0 or 0.5 | n.d.[a] | n.d. | n.a.[b] | n.d. |

[a] not determined. [b] not applicable.

**Supplementary Table 3.** Doubling times

| Strain | Description | | Doubling time (min) | |
|---|---|---|---|---|
| | Plasmid | BzF (mM) | 30 °C | 37 °C |
| DH10β | No plasmid | 0 | 42 ± 6[a] | 30 ± 5 |
| DH10β Δ*cyaA* | No plasmid | 0 | 95 ± 11 | 67 ± 6 |
| DH10β Δ*cyaA* | F1RP-GFP, F2P, SP | 0 | 94 ± 4 | 90 ± 9 |
| DH10β Δ*cyaA* | F1RP-GFP, F2P, SP | 0.5 | 94 ± 6 | 86 ± 8 |
| DH10β Δ*cyaA* | F1RP-GFP, F2P, $SP^{null}$ | 0 | 70 ± 9 | 62 ± 9 |
| DH10β Δ*cyaA* | F1RP-GFP, F2P, $SP^{null}$ | 0.5 | 93 ± 6 | 85 ± 11 |
| DH10β Δ*cyaA* | F1RP-GFP, F2P, $SP^{null}$-Myo | 0 | 77 ± 6 | 59 ± 6 |
| DH10β Δ*cyaA* | F1RP-GFP, F2P, $SP^{null}$-Myo | 0.5 | 97 ± 8 | 87 ± 11 |
| DH10β Δ*cyaA* | F1RP*-GFP, F2P*, SP | 0 | 138 ± 12 | 106 ± 9 |
| DH10β Δ*cyaA* | F1RP*-GFP, F2P*, SP | 0.5 | 88 ± 10 | 91 ± 13 |

[a] n = 4, doubling time precision: 95% confidence interval.

**Supplementary Table 4.** Primers used in this study.

| Primer Name | Primer Sequence (5' to 3') | Description |
|---|---|---|
| oMK-0017 | GGGAATTCCATATGACATCGGAAAACCCGTTACTGG | Cloning of F2P and F2P* |
| oMK-0018 | CGGGGTACCCGGGGATCCTCGAGCAAAGCCTGCTGAGTTAATACGGAATC | Cloning of F2P |
| oMK-0016 | AAAACTGCAGTTAGAGCAAAGCCTGCTGAGTTAATACG | Cloning of F1RP-CAT and F1RP*-CAT |
| oMK-0019 | AAAACTGCAGGGATGACATCGGAAAACCCGTTACTGG | |
| oMK-0056 | GTTGGCGGAATCACAGTCATGACGGGTAGCAAATCAGGCGATACGTCTTGGTGTAGGCTGGAGCTGCTTCGAAG | Amplification of tetracycline resistance gene from pKD4-tet$^R$ for the construction of *cyaA* knockout strain |
| oMK-0057 | CGGATAAGCCTCGCTTTCCGGCACGTTCATCACGAAAAATATTGCTGTAAATGGGAATTAGCCATGGTCCATATGAATATCCTC | |
| oMK-0110 | CGGATAAGCCTCGCTTTCCGGCACGTTCATCACGAAAAATATTGCTGTAAATGGGAATTAGCCATGGTCCATATGAATATCCTC | Cloning of F1RP$^{null}$-CAT |
| oMK-0111 | CGCCAGGTAATCGGTCACCGAATC | |
| oMK-0112 | CGGGGTACCCGGGGATCCTCGAGCAAAGCCTGCTGAGTTAATAC | Cloning of F2P* |
| oMK-0132 | CGGATAACAATTTCACACAGAATTCATTAAAGAGGAG | Cloning of F1RP-GFP, F1RP*-GFP, and F1RP$^{null}$-GFP |
| oMK-0134 | TGATCTAGAGGCCTGTGCTAATGATCAG | |
| oMK-0135 | AATGAATTCTGTGTGAAATTGTTATCCGCTCAC | |
| oMK-0133 | CATTAGCACAGGCCTCTAGATCATTAGTGGTGGTGGTGGTGGTGG | Cloning of F1RP-GFP, F1RP*-GFP, F1RP$^{null}$-GFP, F1RP-GFP-bla, and F1RP-GFP-glnS |
| oMK-0225 | ATGAGCAAAGGAGAAGAACTTTTCACTGGAGTTG | Cloning of F1RP-GFP-bla |
| oMK-0226 | GTGAAGACGAAAGGGCCTCGTGATAC | |
| oMK-0227 | CGAGGCCCTTTCGTCTTCACCGCGGAACCCCTATTTGTTTATTTTTCTAAATACATTC | |
| oMK-0228 | GTGAAAAGTTCTTCTCCTTTGCTCATACTCTTCCTTTTTCAATATTATTGAAGCATTTATCAGGGTTATTG | |
| oMK-0160 | TAATGATCTAGAGGCCTGTGCTAATGATCAG | Cloning of F1RP-GFP-bla and F1RP-GFP-glnS |
| oMK-0213 | AGAAATTACATATGAGCAAAGGAGAAGAACTTTTCAC | Cloning of F1RP-GFP-glnS |
| oMK-0214 | GAAACTGCAGTTAGTGGTGGTGGTGGTGGTGG | |
| oMK-0222 | CAACTCAGCAAAAGTTCGATTTATTCAAGTGAAGACGAAAGGGCCTCGTGATAC | |
| oMK-0223 | TTGAATAAATCGAACTTTTGCTGAGTTGAAGGATC | |
| oMK-0180 | CGAGCAGCTCAGGGTCGAATTTG | Cloning of SP-tRNA$^{null}$ and SP$^{null}$ |
| oMK-0181 | CCTAATGCAGGAGTCGCATAAGGGAG | Cloning of SP-aaRS$^{null}$ and SP$^{null}$ |
| oMK-0186 | GTTTGTGAGCTCCCGGTCATCAATC | Cloning of SP-aaRS$^{null}$ |
| oMK-0200 | GCTCACATGTTGCGAAGCGGAATTAC | Cloning of SP-tRNA$^{null}$ |

| | | |
|---|---|---|
| oMK-0058 | GAATGCCGCGTGTTACCGTTGAT | Verification of DH10β Δ*cyaA* strain construction |
| oMK-0059 | CAGTCAGTTTCCGCTAAGATTGCATGC | |
| oMK-0028 | GTCTGCTTACATAAACAGTAATACAAGGGGTGTT | |
| oMK-0465 | TAAGTCGACGCGTTTAAACGGTCTC | Cloning of $SP^{null}$-Myo |
| oMK-0466 | CATAGATCTGCACCTCCTTTGTGAAATTGTTATC | |
| oMK-0467 | CACAAAGGAGGTGCAGATCTATGGTTCTGTCTGAAGGTGAATGGCAG | |
| oMK-0468 | CCGTTTAAACGCGTCGACTTAACCCTGGTAACCCAGTTCTTTGTATTTAGC | |
| oMK-0530 | GGAGAAGGATTTATGAGTTCTTCAAAAGGGAATTTTATAGCTGTTGATGAC | Cloning of SP-K204F and SP-K204F-$tRNA^{null}$ |
| oMK-0531 | ATCCAAACCCGTTAAGACAGGGTTGTG | |
